# Structural and functional insights into the enzymatic plasticity of the SARS-CoV-2 NiRAN Domain

**DOI:** 10.1101/2023.09.25.558837

**Authors:** Gabriel I. Small, Olga Fedorova, Paul Dominic B. Olinares, Joshua Chandanani, Anoosha Banerjee, Young Joo Choi, Henrik Molina, Brian Chait, Seth A. Darst, Elizabeth A. Campbell

**Affiliations:** Laboratory of Molecular Biophysics, The Rockefeller University, 1230 York Avenue, New York, NY 10065, USA; Department of Chemistry and Department of Molecular, Cellular and Developmental Biology, Yale University, New Haven, CT 06511, USA; Howard Hughes Medical Institute, Yale University, New Haven, CT 06520, USA; Laboratory of Mass Spectrometry and Gaseous Ion Chemistry, The Rockefeller University, New York, NY, USA; Proteomics Resource Center, The Rockefeller University, New York, NY 10065, USA; University of North Carolina at Chapel Hill School of Medicine, Chapel Hill, NC 27599, USA

**Keywords:** NiRAN domain, SARS-CoV-2, capping, mRNA cap, RNAylation, deRNAylation, NMPylation, cryo-EM, coronavirus

## Abstract

The enzymatic activity of the SARS-CoV-2 nidovirus RdRp-associated nucleotidyltransferase (NiRAN) domain is essential for viral propagation, with three distinct activities associated with modification of the nsp9 N-terminus, NMPylation, RNAylation, and deRNAylation/capping via a GDP-polyribonucleotidyltransferase reaction. The latter two activities comprise an unconventional mechanism for initiating viral RNA 5’-cap formation, while the role of NMPylation is unclear. The structural mechanisms for these diverse enzymatic activities have not been properly delineated. Here we determine high-resolution cryo-electron microscopy structures of catalytic intermediates for the NMPylation and deRNAylation/capping reactions, revealing diverse nucleotide binding poses and divalent metal ion coordination sites to promote its repertoire of activities. The deRNAylation/capping structure explains why GDP is a preferred substrate for the capping reaction over GTP. Altogether, these findings enhance our understanding of the promiscuous coronaviral NiRAN domain, a therapeutic target, and provide an accurate structural platform for drug development.

## Introduction

*Nidovirales* are an order of positive-sense, single-stranded RNA viruses that includes the Coronavirus (CoV) family^1^, etiological agents responsible for major and deadly zoonotic events^2,3^. Most recently, SARS-CoV-2 has been responsible for the ongoing COVID-19 pandemic^4,5^, with catastrophic deaths and health, social, and economic impacts worldwide. CoVs possess a large (∼30 kilobases), poly-cistronic RNA genome containing open reading frames (ORFs) for the non-structural proteins (nsps) at the 5’ end, followed by ORFs encoding structural and accessory proteins at the 3’ end^3^ ORF1a and ORF1ab encode two polyproteins, PP1a and PP1ab respectively, that are proteolytically processed into the sixteen nsps responsible for genome expression and replication^3,6^. Central to replication is the conserved RNA-dependent RNA polymerase (RdRp) encoded in nsp12 that, along with essential cofactors nsp7 and two copies of nsp8, comprises the CoV holo-RdRp^7^. Multiple structures of the holo-RdRp with product-template RNA scaffold have been resolved by cryo-electron microscopy (cryo-EM) and are referred to as replication-transcription complexes (RTC)^8^.

The Nidoviral RdRp-associated nucleotidyltransferase (NiRAN) domain is an enigmatic enzymatic activity that is conserved across all *Nidovirales*, unique to the *Nidovirales*, and which is found amino-terminal (N-terminal) to the RdRp on the same nsp (nsp12 of SARS-CoV-2^9^). The NiRAN domain was initially identified in equine arterivirus (EAV) as a Mn^2+^-dependent nucleotidyltransferase. An invariant lysine in the NiRAN domain was the target of selective self-nucleotidyltransferase activity. When conserved residues suggested to be involved in catalysis were mutated, viral replication was largely or completely abrogated ^10^ The CoV NiRAN domain was subsequently demonstrated to contain an orthologous nucleotidyltransferase activity that transferred an NMP moiety to the N-terminal amine of nsp9, a protein without an ortholog outside the CoV family^11^. The NMPylation of nsp9 has been robustly demonstrated *in vitro* across multiple CoVs ^11–14^. However, an *in vivo* role for nsp9 NMPylation has not been uncovered.

Like many eukaryotic mRNAs, CoV RNAs are 5’-capped by a 7-methylguanosine nucleotide linked by a 5’-5’ triphosphate to the 5’-RNA 2’-O-methylated nucleotide^15^. The mRNA caps are crucial for protecting transcripts from degradation^16^ and for translation initiation^17^. Since CoV RNAs are not known to access the infected cell nucleus where the host capping assembly apparatus is located, CoVs encode their own enzymes for generating 5’-caps on the viral RNAs^8^. While nsp14 and nsp16 have long been known to harbor the N7-methyltranserase and 2’-O-methyltransferase activities, respectively^18,19^, the identity of the enzyme that initially generates the 5’-G linkage was only recently confirmed^14^. In a unique two-step mechanism, the NiRAN domain catalyzes the transfer of a 5’-triphosphorylated RNA (pppRNA) to the N-terminus of nsp9, forming a covalent RNA-nsp9 intermediated in a process termed RNAylation. The NiRAN then transfers the RNA from nsp9 (deRNAylation) to GDP, generating the core cap structure GpppN-RNA^14^.

To gain insight into the remarkable enzymatic plasticity of the NiRAN domain, we structurally and functionally characterized two of the three known NiRAN enzymatic activities, UMPylation and deRNAylation/capping. We used a UTP analog with a nonhydrolyzable α-β-phosphate bond (UMPCPP) to visualize a structure of the nsp9-RTC poised for UMPylation of the nsp9 N-terminus at 2.9 Å nominal resolution (∼2.8 Å local resolution in the NiRAN active site). We show that UMPylation of the nsp9-N-terminus may serve to protect cytosolic nsp9 from targeted proteolytic degradation^20^, suggesting a potential *in vivo* role for nsp9 UMPylation. We used a slowly reacting GDP analog (GDP-β-S) to visualize structures of the RNA-nsp9-RTC poised for deRNAylation and GDP transfer at 3.0 Å nominal resolution (∼2.8 Å local resolution in the NiRAN active site). Our results illustrate why GDP is a strongly preferred substrate for the NiRAN GDP-transferase activity^14^. The high local resolution of our structures enables near atomic-resolution insights into the diverse NiRAN catalytic activities and their mechanisms.

### Structure/function analysis of NiRAN-mediated NMPylation of nsp9

The SARS-CoV-2 NiRAN domain efficiently NMPylates the N-terminus of nsp9 *in vitro*^11–14^; in our hands, UMPylation was essentially 100% efficient as monitored by native mass spectrometry (Figure 1A), while NMPylation with other NTPs was less efficient. We used a UTP analog with a nonhydrolyzable α-β-phosphate bond {uridine-5’-[(α,β)-methyleno]triphosphate (UMPCPP)} to capture a pre-catalytic intermediate of NiRAN-mediate nsp9 UMPylation [nsp9-RTC(UMPCPP)] and determined a cryo-EM structure to 2.9 Å nominal resolution (Figure 1B, Figure S1, Table S1). We call this complex UMP-i, for UMPylation intermediate. In UMP-i, nsp9 docks into the NiRAN domain with the conserved nsp9-GxxxG helix (Figure S1F), binding a conserved hydrophobic patch on nsp12. The five N-terminal residues of nsp9 extend through a deep groove on the NiRAN domain surface, terminating with the nsp9 N-terminal Asn (N1) deep in the NiRAN active site (Figures 1C and 1D). The eponymous glycines of the nsp9 GxxxG helix, G100 and G104, are two of six residues in the helix to form a series of hydrophobic interactions with the NiRAN domain.

**Figure 1.**
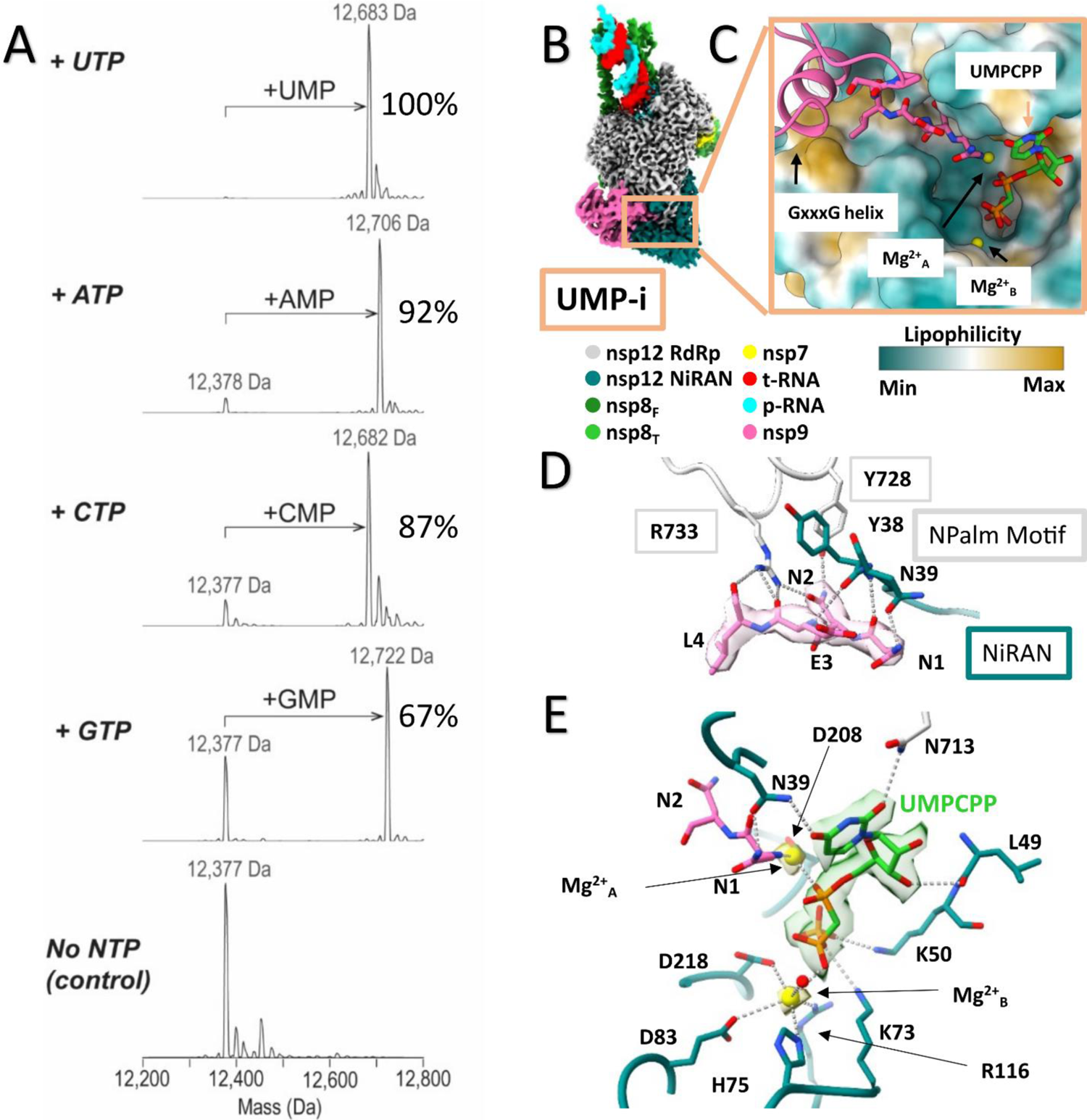
Structural and biochemical characterization of nsp9 NMPylation. (A) Deconvolved native MS spectra showing the mass shift corresponding to the mono-NMPylation of nsp9 from incubation of nsp9 with WT nsp12 and four different NTPs. The measured masses were within 1 Da relative to the expected masses. Efficiency of NMPylation shown beside each panel. (B) Locally filtered cryo-EM map of the 2.8 Å nominal resolution structure reveals the general architecture of an NMPylation intermediate, termed UMP-i here. Map is colored according to fitted model chains. The two copies of nsp8 are noted as nsp8-thumb (nsp8T) and nsp8-finger (nsp8F) (C) Close up of the NiRAN domain active site. Nsp12 is portrayed as a surface and is colored by lipophilicity. (D) Close up of the interface between nsp9’s N-terminal extension and nsp12. Interacting residues are shown as sticks. Nsp9 is shown within locally filtered cryo-EM map and polar interactions are shown with dashed gray lines. (E) Close up of the NiRAN domain active site showing the structural basis of ligand binding for NMPylation. Interacting residues are shown as sticks. UMPCPP and the Mg^2+^ ions are shown within locally filtered cryo-EM map and polar interactions are shown with dashed gray lines.

Nsp9 can dimerize in solution, and nsp9 crystallizes as a dimer with the GxxxG helix at the dimer interface^21,22^. Mutations in conserved residues of the nsp9 GxxxG helix disrupt nsp9 dimerization in solution, and these same mutations result in loss of viral fitness^23,24^. From these results it was concluded that nsp9 dimerization was required for viral fitness, however it is possible that nsp9 dimerization is an *in vitro* artifact and the loss in fitness is due to impaired ability to bind nsp12.

The N-terminal loop of nsp9 extending into the NiRAN domain active site comprises five conserved residues, most notably the invariant N-terminal tripeptide NNE (Figure S1F). The nsp9 N-terminus is positioned for catalysis by a hydrogen bond (H-bond) between the sidechain of nsp9 N1 and the sidechain of nsp12 N39, and the interface continues with a H-bonding network between nsp9 N2, E3 and L4 and nsp12 Y38, Y728, and R733 (Figure 1D, Figure S1G). The role of the nsp12 palm domain residues in NiRAN domain functions has previously been noted^14,25^, and the residues are conserved within CoVs (Figure S1G). Due to the importance of this motif in facilitating nsp9 binding and other NiRAN domain functions, we call these conserved nsp9-palm residues the **<u>N</u>**iRAN-associated **<u>Palm</u>** (NPalm) motif. Despite the absolute conservation of nsp9 E3 (Figure S1F), the sidechain does not appear to play a role in binding nsp12, suggesting a potential role in other nsp9 functions.

Within the NiRAN domain active site pocket, the UMPCPP is positioned in a ‘base-up’ pose, distinct from previously observed ‘base-out’ or ‘base-in’ NiRAN NTP binding poses^26^, with its α-phosphate positioned for catalysis (Figure 1E). Structural modeling indicates that base-in binding of GTP in the NiRAN G-site^26^ would sterically clash with the base-up NMPylation pose, explaining the lower efficiency of GMPylation (Figure 1A, Figure S1H). The UMPCPP is positioned by a network of polar interactions, including H-bonds between the nucleobase functional groups O2 and O4 and NPalm N713 and NiRAN-N39, respectively (Figure 1E). The NiRAN active site and UMPCPP substrate coordinate two Mg^2+^ ions important for the NMPylation reaction (Figure 1E). Mg^2+^_A_ is coordinated close to the pending phosphoramidate bond, poised to catalyze the reaction that would break the α-β phosphodiester bond in the natural substrate UTP (Figure 1E). Mg^2+^_B_ is coordinated close to the β- and γ-phosphates, likely stabilizing the PPi leaving group^27^.

### UMPylation protects the nsp9 N-terminus from deamidation by N-terminal asparagine

#### amidohydrolase

In eukaryotic cells, an N-terminal asparagine is a tertiary destabilizing residue in the N-end rule ubiquitin-dependent pathway^20^. The first enzyme in this pathway, N-terminal asparagine amidohydrolase (NTAN1), deamidates the N-terminal asparagine^28^, converting it into aspartate, a substrate for the Arginine-transferase reaction, the target for ubiquitin ligases. This pathway can reduce the half-life of proteins in eukaryotic cells with N-terminal asparagines by over 400-fold^20,29^ The N-terminal asparagine of nsp9 is invariant across CoVs (Figure S1F) and required for efficient NMPylation^11^. Because nsp9 is required for CoV viability, its degradation would interfere with infection and replication ^11,14^. Therefore, we investigated if nsp9 could serve as a substrate for human NTAN1 (hNTAN1) *in vitro*. Unmodified nsp9 was deamidated by hNTAN1, while UMP-nsp9 was completely protected (Figure S1I), suggesting UMPylation could serve as a protectant against hNTAN1 modification.

#### NiRAN domain RNA substrate preferences

In addition to nsp9 NMPylation, the CoV NiRAN domain mediates RNAylation of the nsp9 N-terminus as well as the deRNAylation of nsp9 and transfer of GDP to the RNA to form the 5’-GpppA cap^14^. The NiRAN-mediated RNAylation and deRNAylation reactions are inefficient *in vitro*^14^ compared to NMPylation. We hypothesized that RNAylation and/or deRNAylation may be facilitated by secondary structure in the substrate RNA. To test this hypothesis, we examined three 5’-triphosphorylated model RNAs based on the 5’-UTR of the SARS-CoV-2 genome (Figure S2A), an unstructured 10mer, a 20mer, and a 33mer comprising the entire 5’ UTR through the conserved stem-loop 1^30^. The 20mer RNA is predicted to form a stable hairpin not found in the SARS-CoV-2 genome^30^ but could represent a nascent RNA intermediate structure. Monitoring the time-course of NiRAN-mediated nsp9 RNAylation with each pppRNA revealed a slight preference for the shorter, unstructured 10mer pppRNA (Figures S3B and S3C). To test the effect of RNA secondary structure on the deRNAylation/capping reaction, we purified the three different RNAylated-nsp9 species and evaluated NiRAN-mediated RNA capping efficiencies using GDP as a substrate. The efficiency and rate of the deRNAylation/capping reaction was not affected by RNA length or secondary structure propensity (Figures S3D and S3E).

DeRNAylation of RNA-nsp9 has been previously demonstrated with both GDP and GTP^14,31^. As observed by Park et al. (2022), the NiRAN domain strongly prefers GDP over GTP for 20mer RNA-nsp9 deRNAylation (Figures 2A and 2B). The efficiency of a deRNAylation reaction with GTP was restored by the addition of a stoichiometric amount of the nsp13 helicase (Figures 2A and 2B), indicating that the NTPase activity of nsp13^32^ can convert the GTP pool into mostly GDP for efficient GDP-PRNTase activity. With nMS, we monitored mRNA capping (Figures 2C and 2D), and we observed the formation of the canonical 5’ mRNA cap when either GDP or GTP was used in the reaction (Figure 2C), distinct from the formation of a noncanonical mRNA cap produced in Rhabdoviruses when GTP is utilized as a substrate^33^. Monitoring the reaction with nMS allows us to observe the capping reaction beginning with RNAylation of nsp9 and followed by the deRNAylation of nsp9 and the capping of the mRNA with the preferred substrate, GDP, as well as with GTP (Figure 2D).

**Figure 2.**
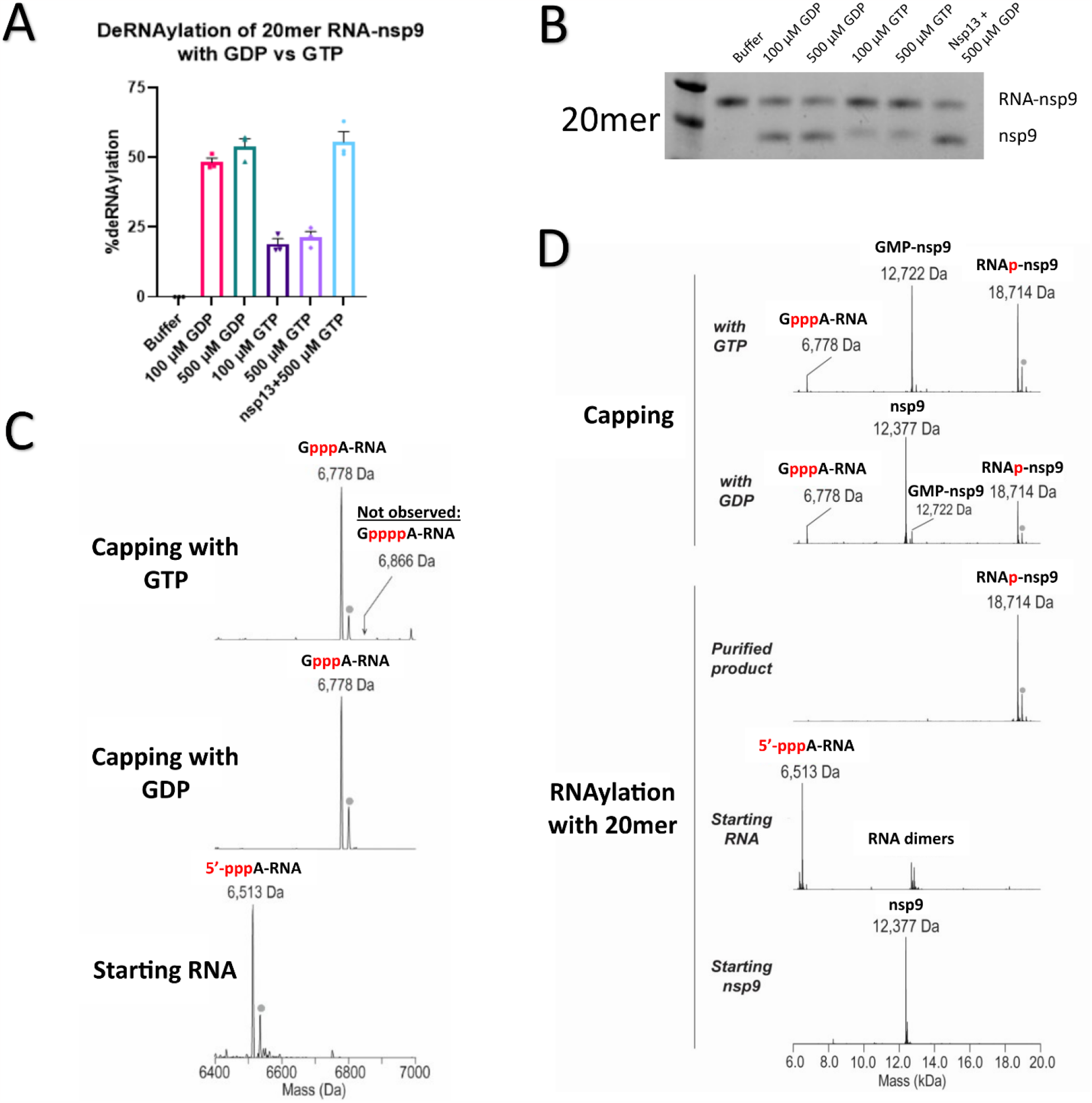
Biochemical analysis of NiRAN domain capping with GDP versus GTP. (A) Comparison of NiRAN domain mediated deRNAylation of 20-mer RNA-nsp9 with GDP, GTP, and GTP+nsp13. Reactions were quenched in 4X LDS + 50 mM DTT, run on SDS-PAGE, and then bands were quantified using gel densitometry. Numbers used for analysis are the intensity of the nsp9 band divided by the sum of the RNA-nsp9 and nsp9 band intensities. These are the results of 3 independent experiments. (B) Representative gel for the deRNAylation experiment shown in (A). (C) nMS analysis of NiRAN domain mediated deRNAylation of nsp9 and resultant GpppA-RNA cap production with GDP and GTP. The gray dot indicates a satellite peak for Mg^2+^ adduction. The position of a hypothetical GppppA peak is noted for reference. (D) Full nMS spectra of those shown in (C) monitoring RNAylation with a 20mer and then capping with GDP or GTP

#### Structural analysis of a NiRAN domain capping intermediate

To understand the structural basis of the NiRAN domain GDP-PRNTase/capping activity, we determined 3.0 Å resolution capping intermediate cryo-EM structures of the RTC in complex with RNAylated-nsp9 (20mer RNA; Figure 3A) with a poorly reactive GDP analog, GDP-βS (Figures S4A and S4B). After initial steps of processing, the consensus map exhibited features indicative of structural heterogeneity. To improve the maps in the area of interest, we constructed a soft mask encompassing the NiRAN domain and nsp9 and used masked classification with signal subtraction^34^ to identify six conformational states (Figure S3C). Two of the states (C00 and C02; Figure S3C) had very poor or no density for nsp9 and were not further processed. C01 contained strong cryo-EM density for RNAylated-nsp9 resolving the entire 20mer length of the nsp9-linked RNA with the predicted stem-loop (Figure S2A), but the GDP-βS substrate was unable to bind (Figure S5). The remaining classes revealed conformational mobility of nsp9 in its binding to nsp12. C03 and C05 were similar to C04 (except for motion in nsp9) but the maps in the NiRAN active site were inferior to C04. Therefore, our further analysis focused on C01 and C04 (Figure S3C).

**Figure 3.**
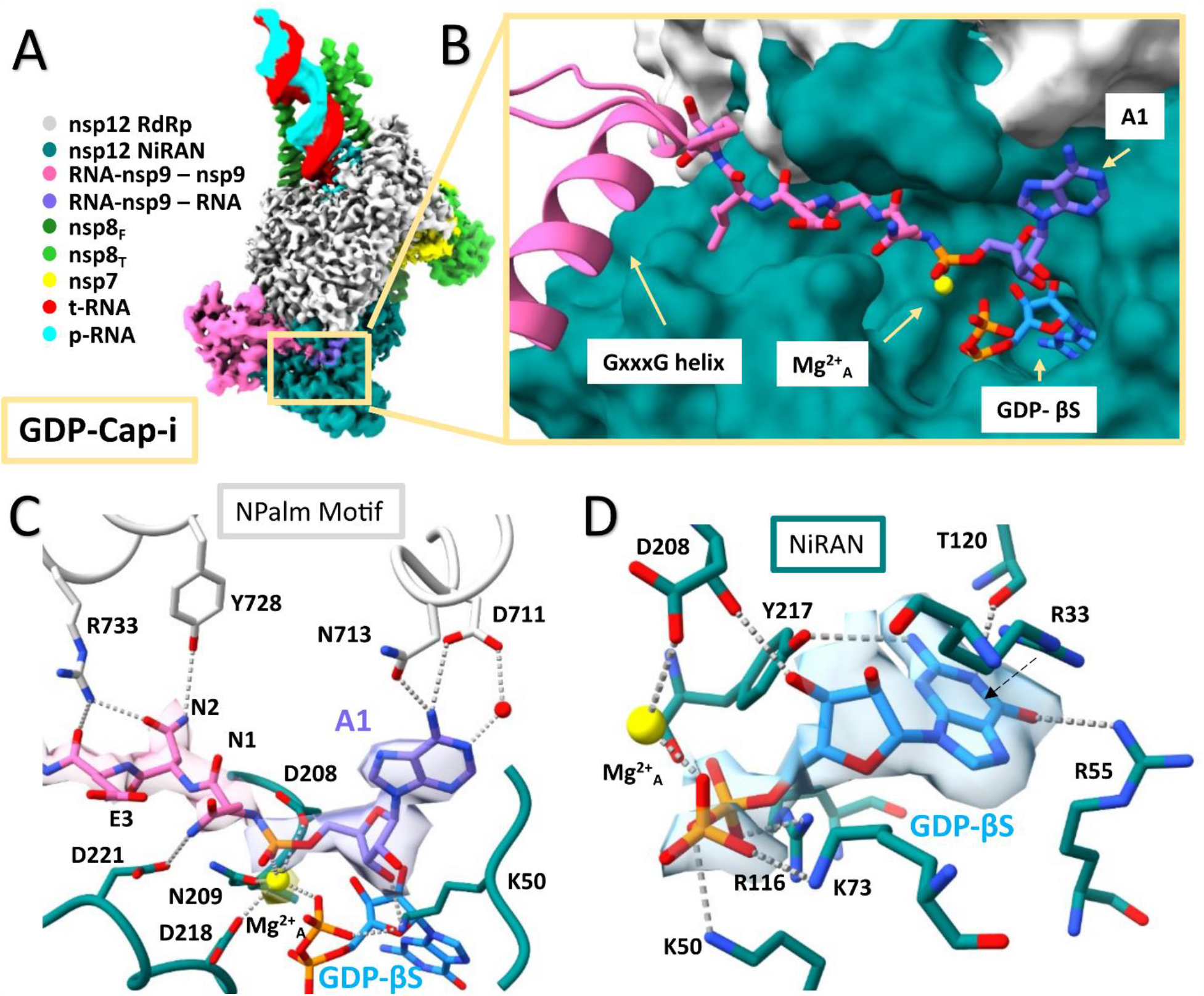
Structural analysis of a NiRAN domain capping intermediate. (A) Locally filtered cryo-EM map of the 3.0 Å resolution structure reveals the general architecture of a trapped catalytic deRNAylation intermediate. Map is colored according to fitted model chains. (B) Close up of the NiRAN domain active site. Nsp12 is portrayed as a surface and is colored according to domain. (C) Close up of the interface between RNA-nsp9’s N-terminal extension and A1 with nsp12. Interacting residues are shown as sticks. RNA-nsp9 is shown within locally filtered cryo-EM map and polar interactions are shown with dashed gray lines. (D) Close up of the NiRAN domain G pocket binding a GDP analog, GDP-βS, and being positioned for catalysis. Interacting residues are shown as sticks and polar interactions are shown with dashed gray lines. The pi-cation interaction is shown with a dashed arrow. GDP-βS is shown within a locally filtered cryo-EM map.

The C04 class contained strong cryo-EM density for RNAylated-nsp9 and the GDP-βS (we designate this structure GDP-Cap-i; Figures 3A and S3C, Table S1). The nominal resolution of the structure was 3.0 Å, with local features in the NiRAN active site resolved to 2.8-2.9 Å (Figure S4F). In GDP-Cap-i, RNA-nsp9 docks into the NiRAN domain with the same interface as the UMP-i structure, with the GxxxG helix forming an extensive interface with nsp12 and the N-terminus of nsp9 loaded into the NiRAN active site (Figures 3A and 3B). The first nucleotide of the nsp9-linked RNA (A1) was well resolved, binding in the same base-up pose as the UMPCPP of UMP-i (Figures 1C and 3B). Like the UMPCPP, the A1 base interacts with NPalm residue N713 and forms a direct and a water-mediated interaction with D711 (Figure 3C). Deeper in the NiRAN active site, the GDP-βS binds in the G-site in the base-in pose (Figure 3D), with the guanine base recognized by the same conserved NiRAN residues an in the GTP-bound NiRAN^26^. We modeled a Mg^2+^_A_ in density located between the β-phosphate of the GDP-βS and the phosphoramidate bond of RNA-nsp9 (Figures 3B-D), coordinated by the phosphoramidate phosphate, NiRAN residues D208, N209, and D218, and the GDP-βS β-phosphate (Figure 3C). It is likely critical for catalyzing the deRNAylation/capping activity. There were no sites with the appropriate cryo-EM density and coordination geometry for a second Mg^2+^-ion, which would normally stabilize the leaving group^27^; leaving group stabilization may not be required for this reaction since the leaving group is the protein nsp9.

The second analyzed class, C01 (nominal resolution 3.1 Å; Figure S3C), contained strong cryo-EM density for RNAylated-nsp9, but here the entire 20mer length of the nsp9-linked RNA, including the predicted stem-loop (Figure S2A) was resolved (Figures S5A and S5B). We term this structure 20mer-Stem-Loop (20mer-SL). The 5’- and 3’-ends of the SL-RNA interact with the surface of the NiRAN and NPalm (Figure S5C). To accommodate the positioning of the SL, the four 5’-nucleotides of the nsp9-linked RNA (A1-U2-U3-A4) are constrained in a sharp turn, causing a shift in the position of the nsp9-linked A1 and the ordering of the U2-phosphate, introducing a severe clash with the β-phosphate of the GDP (modeled from the GDP-Cap-i structure; Figure S5D). Thus, although the nsp9 N-terminus, nsp9-linked A1, the Mg^2+^_A_-ion, and other elements of the NiRAN active site are positioned similarly to the GDP-Cap-i structure (Figures 3C and S5C), GDP is unable to bind, rendering the 20mer-SL state catalytically inactive. Thus, the catalytically active GDP-Cap-i and inactive 20mer-SL states are in equilibrium. Surprisingly, the deRNAylation/capping reaction is not measurably slower with this 20mer-SL RNA compared with the unstructured 10mer (Figure S2D), suggesting that the folding/unfolding transition of the RNA-SL is not rate limiting.

#### Chemical mechanism of NiRAN domain catalysis

The NiRAN domain enzymatic activities have been well studied^10–12,14^. Here, we build on these studies by resolving two distinct pre-catalytic intermediates of NiRAN-mediated reactions at high-resolution (< 3 Å in the NiRAN active site; Extended Data Figures S1C and S5F). This allows for an atomic-level structural understanding of the chemical mechanism underlying two NiRAN activities (Figure 4), providing a structure-based platform for therapeutics targeting this essential activity in CoVs.

**Figure 4.**
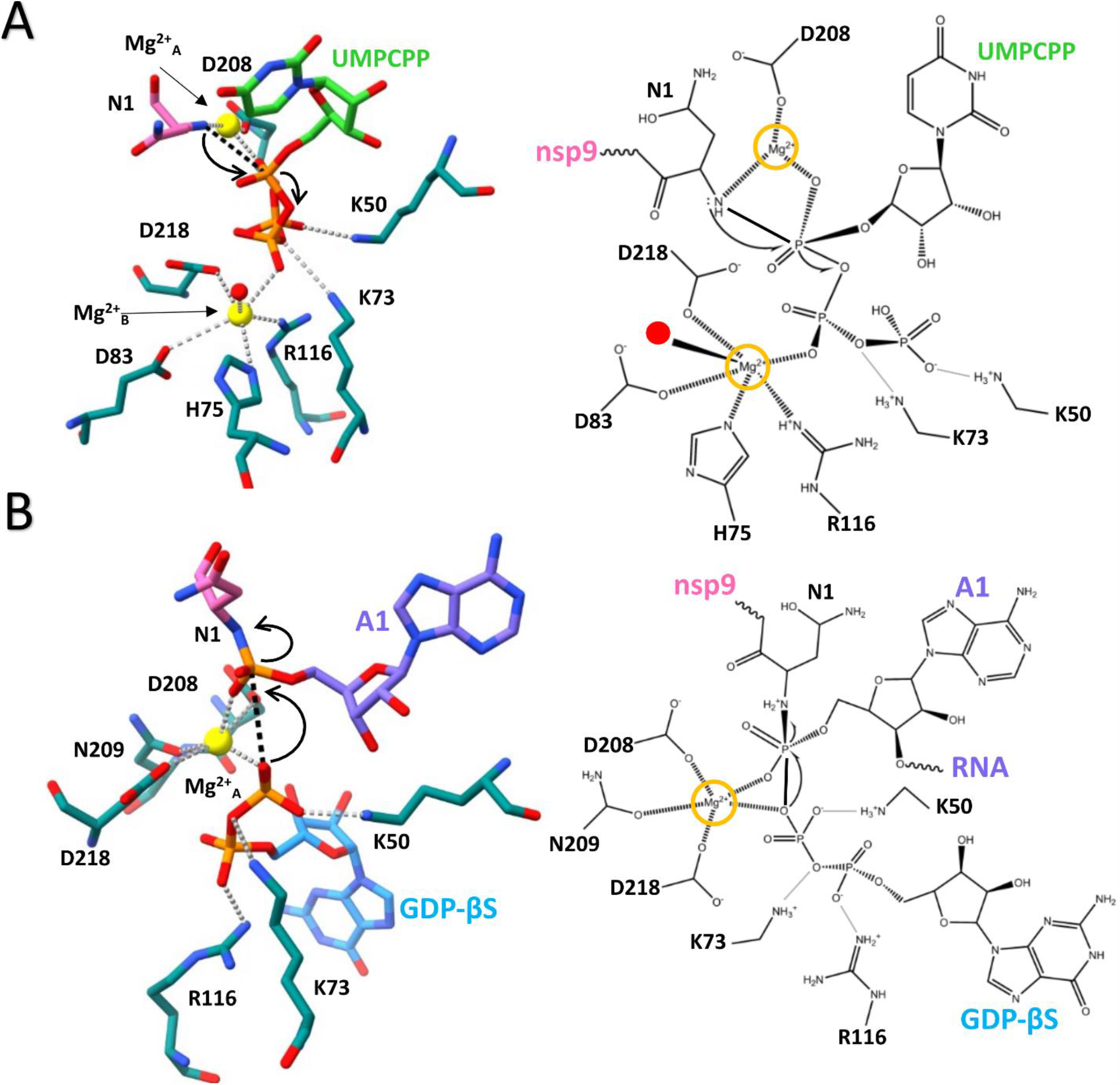
Chemical reaction schema for NMPylation and RNA capping. (A) Reaction schema for NiRAN domain mediated NMPylation of nsp9. Right panel is a 2D visualization of the left panel. Polar interactions shown with dashed gray lines. New bond is shown with black dashed line and arrows show direction of electron transfer. (B) Reaction schema for NiRAN domain mediated RNA capping. Right panel is a 2D visualization of the left panel. Polar interactions shown with dashed gray lines. New bond is shown with black dashed line and arrows show the directions of electron transfer.

In NiRAN-mediated NMPylation of nsp9, the N-terminal amine of nsp9 forms a phosphoramidate bond with the α-phosphate of the incoming NTP substrate, releasing PPi^11,12,14^. In the NMPylation intermediate (UMP-i), the N-terminal amine of nsp9 is positioned to execute an S_N_2 nucleophilic attack on the α-phosphate of the nucleotide substrate bound in the base-up pose, catalyzed by Mg^2+^_A_ (Figure 4A). Completion of the reaction results in the formation of a phosphoramidate bond between the nsp9 N-terminus and the NTP substrate α-phosphate, hydrolysis of the α-β phosphodiester bond, and a PPi leaving group stabilized by Mg^2+^_B_ (Figure 4A). This is reminiscent of the two-metal ion mechanism for nucleic acid polymerases^27^.

In NiRAN domain mediated RNA deRNAylation/capping, the phosphoramidate bond between an RNA chain and the nsp9 N-terminal amine is broken by a GDP, resulting in a canonical 5’-RNA GpppA cap (Figure 2C)^14^. In the RNA capping intermediate GDP-Cap-i, the β-phosphate is positioned to execute an S_N_2 nucleophilic attack on the phosphoramidate phosphorus, catalyzed by Mg^2+^ _A_(Figure 4B). This results in the formation of the GpppA cap, a broken phosphoramidate bond between nsp9 and the RNA, and the recycling of the nsp9 N-terminal amine to an unmodified state. As previously mentioned, the second Mg^2+^-ion (Mg^2+^_B_) for the deRNAylation/capping reaction is not present since the capping reaction does not produce a highly negatively charged PPi leaving group.

In both the UMP-i and GDP-CAP-i pre-catalytic intermediates, the same two NiRAN Lys residues, K50 and K73, are positioned to play potentially important catalytic roles (Figure 4). In the initial discovery of NiRAN nucleotidylation activity, the EAV NiRAN domain was shown to self-NMPylate, covalently attaching an NMP moiety through a phosphoramidate bond to K94 (corresponding to SARS-CoV-2 nsp12 K73^10^). SARS-CoV-2 K73/EAV K94 is absolutely conserved in Nidovirus NiRAN domains^10^ but in the UMP-i structure, K73 is positioned to interact with the UMPCPP β-phosphate and is not close to the α-phosphate, where it would need to be to become covalently linked (Figure 4A). Although many of the NiRAN active site residues are absolutely conserved among all Nidoviruses, the NiRAN domain overall is divergent in sequence; for example, the SARS-CoV-2 and EAV NiRAN domains are only about 10% identical in sequence (among 135 aligned residues). Moreover, Arteriviruses such as EAV do not encode an ortholog to CoV nsp9, so the NiRAN-mediated mechanisms for nucleotidylation and/or capping must be very different than CoVs. SARS-CoV-2 K50 is absolutely conserved among CoVs but is an arginine in Roniviruses and Arteriviruses (such as EAV). Arginines in enzyme active sites typically play electrostatic stabilization roles rather than catalytic roles due to the poor reactivity of Arg^35^. On this basis, we propose that SARS-CoV-2 nsp12 K50 provides a nearby positive charge to stabilize the negatively charged transition state, while K73 plays a more direct role in catalysis, such as a general acid^36^.

### The NiRAN domain accommodates multiple nucleotide binding poses

The NiRAN domain has previously been characterized to bind GTP in a base-in binding mode, while binding other nucleotides in a non-specific base-out mode^13,25,26,37^. In this study, we identified a distinct base-up binding pose for UMPCPP and A1 in the UMP-i (Figure 1) and GDP-Cap-i (Figure 3) structures, respectively (Figure 5A). The UMPCPP (UMP-i) and A1 (GDP-CAP-i) bind similarly, interact with the same loop from the NPalm motif (Figure 5A), and are catalytic intermediates relevant to the NiRAN-mediated NMPylation and capping activities, respectively. The base-in pose (Figure 5A) is specific for guanosine nucleotides (GDP; GDP-CAP-i, or GTP)^13,38^, which are used by the NiRAN domain as capping substrate. The previously described base-out binding mode^25,37^ is likely not relevant to any catalytic activity.

**Figure 5.**
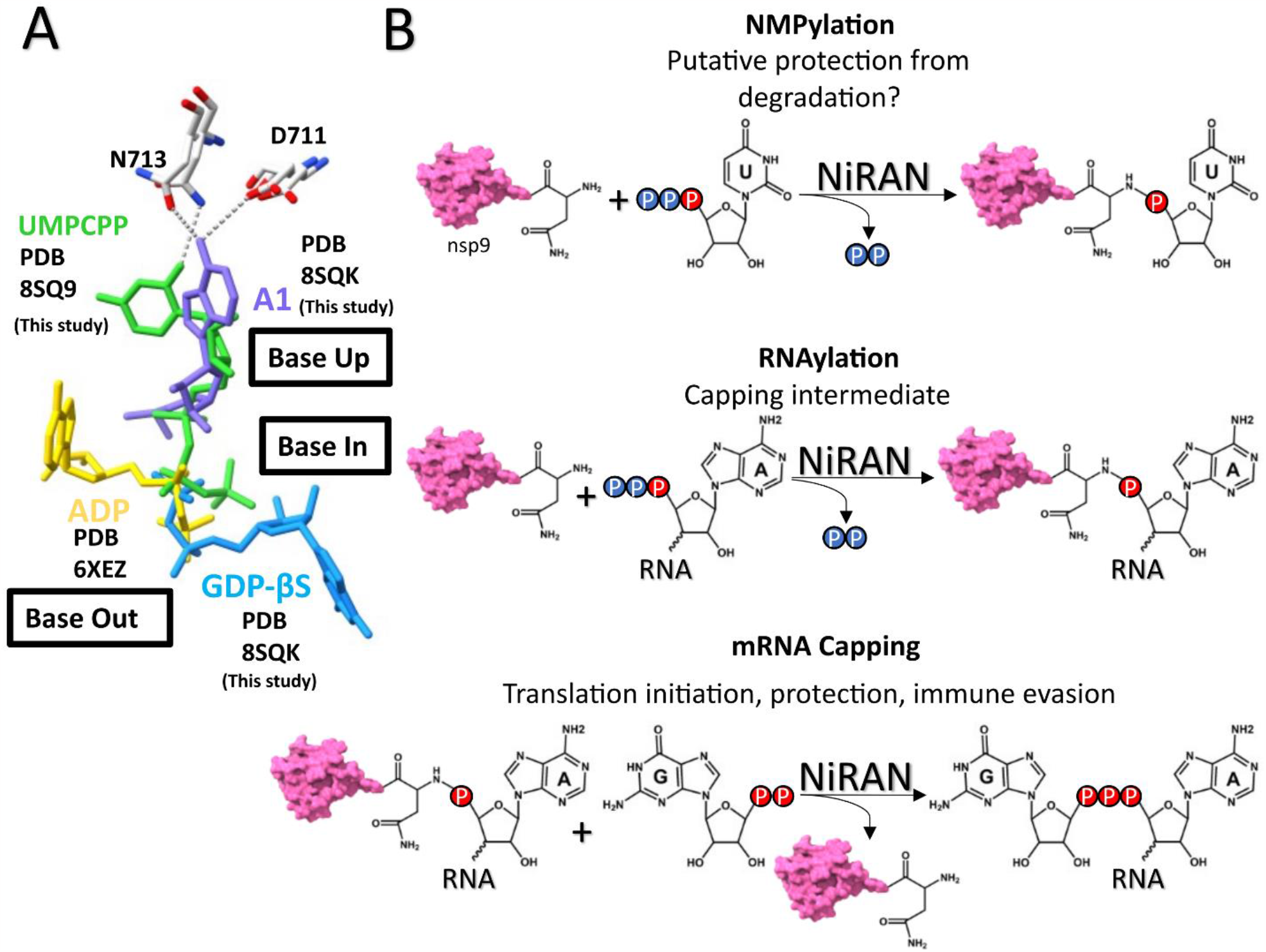
The NiRAN domain binds nucleotides in diverse poses allowing for diverse catalytic activities. (A) Four different structures of the NiRAN domain with nucleotides showcasing the three documented NiRAN domain nucleotide binding poses. Two NPalm motif residues that are critical for promiscuous nucleobase recognition shown in sticks. Polar interactions are shown with dashed gray lines. (B) Schematic of the NiRAN domain’s three documented activities. Proposed role of NMPylation and the roles of RNAylation and mRNA capping are listed beneath the name of the activities.

## Discussion

NiRAN domain function is essential for CoV propagation^10,11^. The NiRAN was initially characterized as a nucleotidyltransferase that, in EAV, NMpylates itself^10^, or in CoVs NMpylates the N-terminus of nsp9. Subsequently, the NiRAN was shown to mediate a remarkable series of reactions, RNAylation of the nsp9 N-terminus, and deRNAylation/capping, that are critical first steps for the essential process of viral 5’-RNA capping^14^. Our detailed structural characterization of catalytic intermediates for two of these three known SARS-CoV-2 NiRAN domain enzymatic activities (Figure 5B), NMPylation (Figure 1) and deRNAylation/capping (Figure 3), provides molecular insight into how the NiRAN domain can mediate so many distinct enzymatic activities. Our results provide structural insight into NMPylation and RNA capping intermediates, revealing the determinants of substrate binding and the chemistry catalyzed by the enzyme, plus providing a platform for structure-based drug development.

### The NiRAN-mediated UMPylation intermediate reveals a new NTP binding pose

In our UMP-i stucture we reveal a new base-up NTP binding pose in the NiRAN domain. Therefore, there are now three distinct base binding poses: 1) base-out, a nonspecific pose that relies on phosphate interactions^25,37^ and is unlikely to have catalytic relevance; 2) base-in, specific for guanosine nucleotides used as the 5’-RNA capping substrate^13,38^ (Figures 3D and 5A); and 3) base-up, a flexible nucleobase binding mode for UMPylation (UMP-i structure; Figures 1B-E) and for the first adenine base of the nsp9-linked RNA (GDP-CAP-i structure; Figures 3B and 3C). We determined the base-up pose for the non-hydrolyzable UTP analog UMPCPP, but we propose that all NMPylation proceeds through each NTP binding in a similar base-up pose seen for UMPCPP (Figures 1C and 1E).

### The role of NiRAN domain mediated NMPylation of nsp9

Nsp9 has been proposed as a target for broad-spectrum antivirals targeting CoVs^31,39–41^, and therefore understanding nsp9 functions, location, and stability is critical. Nsp9, with its absolutely conserved N-terminal Asn (Figure S1F) is a likely target for the N-end rule pathway, which can reduce the cellular half-life of proteins with Asn at the N-terminus by over 400-fold^20,29^. We show that the nsp9 N-terminal Asn can serve as a substrate for hNTAN1 (Figure S1I), the first enzyme involved in this pathway^20,28,29^. Given the reversibility of NMPylation^12,14^, we propose that a possible role for nsp9 NMPylation in SARS-CoV-2 pathogenesis could be to protect nsp9 from degradation, maintaining high nsp9 levels to be used as a critical RNA capping intermediary.

### Substrate preference for NiRAN-mediated deRNAylation/capping

GDP is the preferred substrate over GTP for deRNAylation/capping^14^ (Figures 2A and 2B). Superimposing the G-site GTP from 7UOB^26^ onto the GDP-CAP-i structure reveals that the GTP γ-phosphate closely approaches the nsp9-linked 5’-terminal phosphate of the RNA (2.4 Å closest approach), creating strong electrostatic repulsion in the absence of a neutralizing divalent cation (Figure S4I), explaining the GDP preference. The structure of a catalytic intermediate with a non-hydrolyzable GTP analog would explain how GTP can function as a (non-preferred) capping substrate^14^ (Figures 2A and 2B) and provide further insight into NiRAN domain enzymatic plasticity. The preference for GDP is difficult to understand since the cellular concentration of GTP is expected to greatly exceed that of GDP^42^. We show that the NTPase activity of nsp13 can supply GDP for the deRNAylation/capping reaction (Figures 2A and 2B). The structure of a deRNAylation/capping intermediate with non-hydrolyzable GTP analog GMPPNP was recently reported31 (PDB: 8GWE, EMDB: 34310) ^31^. However, this structure lacks the density necessary for atomic modeling of the GTP analog and metals and does not provide insight into the capping mechanism (Figure S4J).

### NiRAN-mediated RNAylation

Questions remain regarding the NiRAN-mediated RNAylation reaction (Figure 5B). The NiRAN-mediated NMPylation and capping reactions are promiscuous to multiple divalent cations, particularly Mn^2+^ and Mg^11,12,14^, but the RNAylation reaction appears to strictly require Mn^11,14^. Moreover, the NiRAN-mediated RNAylation strongly prefers an A at the RNA 5’-end^14^. The 5’-A of the RNAylation substrate may bind in the same base-up pose as A1 of the nsp9-linked RNA in the GDP-CAP-i structure. A structure of an RNAylation intermediate will be required to confirm this supposition, to understand the unique divalent cation requirement of the RNAylation reaction, and to provide further insight into the NiRAN domain enzymatic plasticity.

### Conclusion

In summary, we have provided the structural basis for the NiRAN domain mediated NMPylation of nsp9 as well as proposed a function for the activity in extending the half-life of nsp9. Importantly, we provide the structural basis for the use of GDP as a substrate for deRNAylation/capping. Altogether, these studies provide insight into the binding determinants and poses of the substrates required for two of the three NiRAN-mediated enzymatic activities. Our results demonstrate the determinants of binding and the poses of the substrates to this enigmatic domain. Our results elucidate critical CoV biology while providing a high-resolution platform for structure-based therapeutic targeting and development against the essential SARS-CoV-2 NiRAN domain.

## Methods

### Expression and purification of nsp7/8

SARS-CoV-2 nsp7/8 was expressed and purified as previously reported^37^. In brief, a pCDFDuet-1 plasmid containing His_6_-PPX-nsp7/8 (Addgene: 159092) was transformed into *E. coli* BL21(DE3) (Agilent) and plated on LB-agar containing 50 μg/mL streptomycin. Cells were grown in LB media supplemented with 10 μM ZnCl_2_, grown to OD_600_ = 0.6 at 30°C at 200 rpm, induced with 0.1 mM IPTG, then incubated for 14 hours at 16°C. Cells were collected via centrifugation, resuspended (in 20 mM Tris-HCl pH 8.0, 300 mM NaCl, 0.1 mM EDTA-NaOH pH 8.0, 5 mM imidazole, 5% glycerol (v/v), 10 μM ZnCl_2_, 1 mM BME, 1x Protease Inhibitor Cocktail (Roche), 1 mM PMSF), and lysed through a French Press (Avestin). The cleared lysate from centrifugation was loaded on a HisTrap HP column (Cytiva), washed, and eluted. The eluate was dialyzed overnight with Precission Protease to cleave the His_6_-tag. The cleaved proteins were passed through another HisTrap HP column, and the resulting flow-through was injected onto a Superdex 75 Hiload 16/600 (Cytiva) for size-exclusion chromatography. Glycerol was added to purified nsp7/8 to reach 20% final concentration, aliquoted, flash-frozen with liquid N_2_, and stored at -80°C until use.

### Expression and purification of nsp9

SARS-CoV-2 nsp9 was produced with a physiological N-terminus using a synthetic gene described previously^41^. The plasmid was transformed into BL21 (DE3) cells and plated on LB-agar containing 50 μg/mL kanamycin and grown in LB media to OD_600_ = 0.6 at 37°C at 200 rpm, induced with 0.5 mM IPTG, then incubated for 4 hours at 30°C. Cells were collected via centrifugation, resuspended (in 20 mM HEPES pH 8.0, 300 mM NaCl, 20 mM imidazole, 5% glycerol (v/v), 1 mM BME, 1x Protease Inhibitor Cocktail (Roche), 1 mM PMSF), and lysed through a French Press (Avestin). The cleared lysate from centrifugation was loaded on a HisTrap HP column (Cytiva), washed, and eluted in 20 mM HEPES pH 8.0, 300 mM NaCl, 250 mM imidazole, 5% glycerol, and 1 mM BME. The eluate was dialyzed overnight with Ulp1 to cleave the His_6_-affinity tag. The cleaved protein was injected onto a Superdex 75 Hiload 16/600 (Cytiva) for size-exclusion chromatography and fractions of interest were passed through another HisTrap HP column to isolate tag-free proteins. Glycerol was added to the isolated protein to reach 20% final concentration, aliquoted, flash-frozen with liquid N_2_, and stored at -80°C until use.

### Expression and purification of nsp12

SARS-CoV-2 nsp12 was expressed^43^ and purified^26^ as previously reported. In brief, a pQE-30/pcl-ts ind+ plasmid containing a His6-small ubiquitin-like modifier (SUMO) SARS-CoV-2 nsp12 and untagged nsp7 and 8 (Addgene no. 160540) was transformed into Escherichia coli BL21 cells (Agilent). Cells were grown and protein expression was induced by the addition of 0.2 mM isopropyl β-D-1-thiogalactopyranoside (IPTG), 10 ng ml−1 tetracycline and 50 μg ml−1 nalidixic acid. Cells were collected by centrifugation, resuspended in 20 mM HEPES pH 8.0, 100 mM NaCl, 5% glycerol (v/v), 1 mM DTT, and lysed in a French press (Avestin). The lysate was cleared by centrifugation and purified on a HiTrap Heparin HP column (Cytiva). The fractions containing nsp12 were loaded onto a HisTrap HP column (Cytiva) for further purification. Eluted nsp12 was dialysed, cleaved with His6-Ulp1 SUMO protease, and passed through a HisTrap HP column to remove the SUMO protease. Flow-through was collected, concentrated by centrifugal filtration (Amicon) and loaded on a Superdex 200 Hiload 16/600 (Cytiva). Glycerol was added to the purified nsp12 to a final concentration of 20%, aliquoted, flash-frozen with liquid N2 and stored at −80 °C.

### Expression and purification nsp13

SARS-CoV-2 nsp13 was expressed and purified as previously reported^37^. In brief, a pet28 plasmid expressing His6-nsp13 was transformed into E. coli Rosetta (DE3) cells (Novagen) and plated on LB-agar containing 50 μg/mL KAN and 25 μg/mL CAM. Single colonies were used to inoculate liquid LB/KAN/CAM cultures. Cells were grown at 37°C, induced at 0.6 OD600 by the addition of IPTG (0.2 mM final), then incubated for 17 hr at 16°C. Cells were collected by centrifugation, resuspended in 50 mM HEPES-NaOH, pH 8.0, 500 mM NaCl, 5 mM MgCl2, 5% (v/v) glycerol, 20 mM imidazole, 5 mM BME, 1 mM ATP, 1 mM PMSF and lysed in a French press (Avestin). The lysate was cleared by centrifugation then purified on a HisTrap HP column. Eluted nsp13 was dialyzed overnight into 50 mM HEPES-NaOH pH 8.0, 500 mM NaCl, 5 mM MgCl2, 5% (v/v) glycerol, 20 mM imidazole, 5 mM BME in the presence of His6-Prescission Protease to cleave the His6-tag. Cleaved nsp13 was passed through a HisTrap HP column and the flow-through was collected, concentrated by centrifugal filtration (Amicon), and loaded onto a Superdex 200 Hiload 16/600 (GE Healthcare) in 20 mM HEPES pH 8.0, 500 mM NaCl, 5 mM MgCl2, 5% glycerol, 1 mM DTT). Glycerol was added to purified nsp13 to reach 20% final, aliquoted, and flash-frozen with liquid N2, and stored at -80°C until use.

### Expression and purification of hNTAN1

Human NTAN1 was expressed and purified as previously reported^28^. In brief, the plasmid, pNTAN2FLAGSII, was transformed into BL21 (DE3) cells and plated on LB-agar containing 50 μg/mL kanamycin and grown in LB media to OD_600_ = 0.6 at 37°C at 200 rpm, induced with 1 mM IPTG, then incubated for 16 hours at 25°C. Cells were collected via centrifugation, resuspended in 100 mM Tris-HCl pH 7.5, 150 mM NaCl, 1 mM EDTA, and lysed in a French press (Avestin). The lysate was cleared by centrifugation and applied to a 5 mL column (Kontes) packed with strep-tactin superflow high capacity resin^44^. This was followed by purification with anti-FLAG M2 magnetic beads and elution using 100 μg/mL FLAG peptide then overnight dialysis into 50 mM Tris-HCl pH 7.5, 150 mM NaCl, 1 mM DTT. The protein was used fresh the following day.

### Native mass spectrometry experiments

#### NMPylation sample prep

Nsp9 (10 μM) was incubated with nsp12 (0.05 μM) and one of four NTPs (200 μM) for 1.5 hours at 37°C in 50 mM HEPES pH 8, 15 mM KAc, 2 mM MgCl2, 1 mM DTT before flash freezing the samples.

#### Capping sample prep

RNA-nsp9 constructs (4 μM) were incubated with nsp12 (0.5 μM) and either GDP or GTP (500 μM) at 37°C in 50 mM HEPES pH 8, 15 mM KAc, 2 mM MgCl2, 1 mM DTT for 4 hours before flash freezing the samples.

#### Native Mass Spectrometry (nMS) analysis

The samples were then buffer exchanged into either 300 mM or 500 mM ammonium acetate solution for the nucleotidylation or deRNAylation samples respectively using Zeba microspin desalting columns with a 40-kDa MWCO (Thermo Scientific). The nMS solutions contained 0.01% Tween-20^45^. The samples concentrations used ranged from 2 – 4 µM. For nMS analysis, 2 – 3 µL of each sample was loaded into a gold-coated quartz capillary tip that was prepared in-house and then electrosprayed into an Exactive Plus with extended mass range (EMR) instrument (Thermo Fisher Scientific) with a static direct infusion nanospray source^45^. The MS parameters used include: spray voltage, 1.2 kV; capillary temperature, 150 – 200 °C; in-source dissociation, 0 V; S-lens RF level, 200; resolving power, 8,750 at m/z of 200; AGC target, 1 x 106; maximum injection time, 200 ms; number of microscans, 5; injection flatapole, 8 V; interflatapole, 7 V; bent flatapole, 5 V; high energy collision dissociation (HCD), 45 V; ultrahigh vacuum pressure, 5 – 6 × 10−11 mbar; total number of scans, at least 100. Mass calibration in positive EMR mode was performed using cesium iodide.

For data processing, the acquired nMS spectra were visualized using Thermo Xcalibur Qual Browser (v. 4.2.47). MS spectra deconvolution was performed using the software UniDec v. 4.2.0^46,47^. Data processing and spectra deconvolution were performed using UniDec version 4.2.0^46,47^. The UniDec parameters used were m/z range: 1,000 – 4,000; background substraction: subtract curved at 10; mass range: 5,000 – 30,000 Da; sample mass every 1 Da; smooth charge state distribution, on; peak shape function, Gaussian; and Beta softmax function setting, 20. The measured masses were typically off by 1 Da relative to the expected mass. The expected masses for nsp9 and the 20-mer 5’-pppRNA are 12,378 Da and 6,512 Da, respectively.

#### Purification of UMP-nsp9

Nsp9 (29 μM) was incubated with nsp12 (2 μM) and UTP (10 mM) for 30 minutes at 30°C in 50 mM HEPES pH 8, 1 mM MnCl2, 5 mM DTT. UMP-nsp9 was then purified from nsp12 and excess UTP over a Superose 6 Increase 10/300 GL column (Cytiva) in 20 mM HEPES pH 8, 120 mM KAc, 10 mM MgCl2, 2 mM DTT and eluted in 2 peaks that were assessed via native mass spectrometry to confirm modification.

#### Tandem mass spectrometry to assess hNTAN1 mediated deamidation of nsp9 and UMP-nsp9

Nsp9 or UMP-nsp9 (5 μM) were incubated with or without hNTAN1 for 1 hour at 37°C in 50 mM Tris-HCl pH 7.5, 150 mM NaCl, 1 mM DTT then samples were flash frozen. Protein samples were trypsinized (Promega) in 40mM ammonium bicarbonate (FISHER SCIENTIFIC). Half of each samples was acidified (0.05% formic acid) followed by desalting and concentration by micro solid phase extracted (Empore, 3M)^48^. Fractions of each digest were analyzed by LC-MS/MS: 35-minute analytical gradient (1%B to 40%B, A: 0.1% formic acid, B: 80% acetonitrile, 0.1% formic acid). A 12 cm long/75um inner diameter built-in-emitter column was connected to mass spectrometer operated in high resolution/high mass accuracy mode (Orbitrap Fusion LUMOS or Q-Exactive HF, Thermo). Mass spectrometer was operated in a hybrid date dependent acquisition (DDA)/parallel reaction monitoring (PRM) mode^49^. The following doubly charged peptides were targeted: NNELSPVALR; N[ump]NELSPVALR;

N(deam)NELSPVALR; N[ump]N(deam)ELSPVALR and N(deam)N(deam)ELSPVALR, where ‘ump’ is uridine monophosphate (C9H11N2O8P, Δm of +306.025302 Da) and ‘deam’ is deamidation (H-1N-1O, Δm of + 0.984016). Data were searched against a custom database containing NSP9 concatenated to a background proteome. ProteomeDiscoverer/Mascot were used to query the data. Tandem MS spectra were manually validated to assure the position of the deamidated residues. Signals of relevant peptides were extracted using Skyline(-daily) v/22.2.1.425^50^.

#### Synthesis of 5’-triphosphorylated RNA oligonucleotides

5’-triphisphorylated RNA oligonucleotides were synthesized on the MerMade 12 RNA-DNA synthesizer (Bioautomation) as previously described (1). Base and 2’-hydroxyl group deprotection and subsequent purification was carried out essentially as described^51^. Briefly, for base and phosphate group deprotection and removal of the oligonucleotide from the support, the polymer support was transferred to a glass vial and incubated with 4 ml of the mixture of 28-30% aqueous ammonium hydroxide (JT Baker) and 40% aqueous methylamine (MiliporeSigma) (1:1) at 65 oC for 15 min. Then the solution was cooled on ice for 10 min, transferred to a clean vial, incubated at -80oC for 1h and evaporated to dryness using SpeedVac. Then 500 μL of anhydrous ethanol was added and the mixture was evaporated again to eliminate all traces of water. In order to deprotect 2’-hydroxyl groups, 500 μL of 1M tetrabutylammonium fluoride in tetrahydrofuran was added and the mixture was incubated at room temperature for 36h. Then 500 μl of 2M sodium acetate, pH 6.0 was added and the resulting mixture was evaporated in the SpeedVac until the volume was reduced by half. The mixture was then extracted three times with 800 μL of ethyl acetate and evaporated in the SpeedVac for 15 min followed by an overnight precipitation at -20 oC with 1.6 ml of ethanol. Then the oligonucleotides were dissolved in 750 μl of sterile water, desalted using GlenPak 1.0 columns (Glen Research), ethanol-precipitated again, dissolved in 200 μl of the RNA storage buffer (20 mM MOPS, pH 6.5, 1 mM EDTA) and stored at -80 oC.

#### RNAylation of nsp9 time points

Nsp9 (20 μM) was incubated with nsp12/7/8 (2 μM) and a pppRNA oligo (100 μM) at 37°C for for the listed time points from 1 minute to 1 hour in 50 mM HEPES pH 8, 15 mM KAc, 4 mM MnCl2, 2 mM DTT, 1 mM TCEP and 2.5 units/mL of YIPP (NEB). At each time point reaction was quenched with 4X LDS and 50 mM DTT then run on SDS-PAGE. Gels were imaged then analyzed using ImageJ to quantify the reaction efficiencies. Three independent experiments were performed for each pppRNA construct and Prism (GraphPad Software) was used to graph the results.

#### Purification of RNA-nsp9

Nsp9 (65 μM) was incubated with nsp12/7/8 (2 μM) and a pppRNA oligo (130 μM) at 37°C for 3 hours in 50 mM HEPES pH 8, 15 mM KAc, 4 mM MnCl2, 2 mM DTT, 1 mM TCEP and 2.5 units/mL of YIPP (NEB). RNA-nsp9 product was then purified from the protein and leftover RNA components through a 1 mL HiTrap Q Fast Flow column in a buffer of 20 mM HEPES pH 7.5, 1 mM DTT with a salt gradient. Eluted product was concentrated, and buffer exchanged into buffer of 50 mM HEPES pH 8, 15 mM KAc, 1 mM DTT and flash-frozen with liquid N2, and stored at -80°C until use.

#### Comparison of GDP and GTP as substrates for deRNAylation

RNA-nsp9 constructs (4 μM) were incubated with nsp12 (0.5 μM) and either GDP or GTP (100 or 500 μM depending on the reaction) at 37°C in 50 mM HEPES pH 8, 15 mM KAc, 2 mM MgCl2, 1 mM DTT. For the reactions including nsp13 (0.5 μM), nsp13 was added to the reaction prior to the addition of GTP. After 15 minutes, reaction was quenched with 4X LDS and 50 mM DTT then run on SDS-PAGE. Gels were imaged then analyzed using ImageJ to quantify the reaction efficiencies. Three independent experiments were performed for each pppRNA construct and Prism (GraphPad Software) was used to graph the results.

#### DeRNAylation of RNA-nsp9 time points

RNA-nsp9 constructs (4 μM) were incubated with nsp12/7/8 (0.5 μM) and GDP (100 μM) at 37°C for the listed time points from 1 minute to 1 hour in 50 mM HEPES pH 8, 15 mM KAc, 2 mM MgCl2, 1 mM DTT. At each time point reaction was quenched with 4X LDS and 50 mM DTT then run on SDS-PAGE. Gels were imaged then analyzed using ImageJ to quantify the reaction efficiencies. Three independent experiments were performed for each pppRNA construct and Prism (GraphPad Software) was used to graph the results.

#### Cryo-EM Sample Preparation

Cryo-EM samples of SARS-CoV-2 RTC were prepared as previously described^26,37^. In brief, purified nsp12 and nsp7/8 were mixed in a 1:2.5 molar ratio and incubated at room temperature for 20 minutes at 22°C. An annealed RNA scaffold (Horizon Discovery, Ltd) was added to nsp12/7/8 and incubated at 30°C for 30 minutes. The complex was then buffer exchanged into 20 mM HEPES pH 8, 80 mM KAc, 2 mM MgCl2, 2 mM DTT, incubated at 30°C for 30 minutes again, then purified overa Superose 6 Increase 10/300 GL column (Cytiva). The eluted nsp12/7/8/RNA complex was pooled and concentrated by centrifugal filtration (Amicon).

#### Cryo-EM Grid Preparation

### Nucleotidylation Intermediate

Before freezing grids beta-octyl-glucoside detergent (β-OG), nsp9, and UMPCPP (JenaBiosciences) were added to sample for a final buffer condition at time of freezing of 20 mM HEPES-NaOH, pH 8.0, 80 mM K-acetate, 2 mM MgCl2, 2 mM DTT, 0.07% β-OG, 500 μM UpCpp. Final RTC and nsp9 concentrations were 36 and 90 μM respectively. C-flat holey carbon grids (CF-1.2/1.3-4Au, Protochips) were glow-discharged for 20 s prior to the application of 3.5 μL of sample. Using a Vitrobot Mark IV (Thermo Fisher Scientific), grids were blotted and plunge-froze into liquid ethane at 90% chamber humidity at 4C.

### DeRNAylation Intermediate

Before freezing grids beta-octyl-glucoside detergent (β-OG), RNA-nsp9, and GDP-βS (JenaBiosciences) were added to sample for a final buffer condition at time of freezing of 20 mM HEPES-NaOH, pH 8.0, 80 mM K-acetate, 2 mM MgCl2, 2 mM DTT, 0.07% β-OG, 500 μM GDP-βS. Final RTC and RNA-nsp9 concentrations were 24 and 39 μM respectively. C-flat holey carbon grids (CF-1.2/1.3-4Au, Protochips) were glow-discharged for 20 s prior to the application of 3.5 μL of sample. Using a Vitrobot Mark IV (Thermo Fisher Scientific), grids were blotted and plunge-froze into liquid ethane at 90% chamber humidity at 4C.

#### Cryo-EM data acquisition and processing

##### UMP-i

Grids were imaged using a 300 kV Titan Krios (Thermo Fisher Scientific) equipped with a GIF BioQuantum and K3 camera (Gatan). Movies were collected with Leginon^52^ with a pixel size of 1.0825 Å per px (micrograph dimensions of 5,760 × 4,092 px) over a defocus range of 0.7 to 2.5 μm with a 20 eV energy filter slit. Videos were recorded in counting mode (native K3 camera binning 2) with ∼25 e−/ Å^2^/s exposure rate in dose-fractionation mode with intermediate frames recorded every 50 ms over a 2 s exposure (40 frames per micrograph) to give an electron exposure of ∼51 e−/Å^2^. Dose-fractionated videos were gain-normalized, drift-corrected, summed and dose-weighted using MotionCor2^53^. The CTF was estimated for each summed image using the Patch CTF module in cryoSPARC v.3.2.0^54^. Particles were picked and extracted from the dose-weighted images with box size of 256 px using cryoSPARC blob picker and particle extraction. The entire dataset consisted of 21,859 motion-corrected images with 10,257,549 particles. Particles containing the RTC were sorted from junk particles using cryoSPARC heterogeneous refinement using a template of an RTC monomer^26^ and junk classes that pulled out particles containing nsp9, not found in the template (Extended Data Fig. 1a). This was followed by further particle curation using iterative heterogeneous refinement (n=4), resulting in 1,190,409 curated particles. These particles were refined using cryoSPARC local and global CTF refinements as well as non-uniform refinements (Extended Data Fig. 1a). Particles were further processed through 2 rounds of RELION v.3.1 Bayesian polishing^55^. Polished particles were refined using cryoSPARC non-uniform refinements then masked (Extended Data Fig. 1a) around the NiRAN domain and nsp9 then subtracted outside the mask for cryoSPARC local refinements around the NiRAN domain active site. Locally refined maps were combined with the consensus maps into a composite map for each class using PHENIX ‘combine focused maps’ to aid model building^56^. Local resolution calculations were generated using blocres and blocfilt from the Bsoft package^57^ (Extended Data Fig. 1c). Angular distribution of particle orientations (Extended Data Fig. 1b) and directional resolution (Extended Data Fig. 1d), calculated through the 3DFSC package^58^ are shown for the final class. GSFSC, calculated through cryoSPARC, and map-model FSC, calculated with Phenix mTriage, are shown for the final class (Extended Data Fig. 1e).

##### GDP-Cap-I and 20mer-SL

Grids were imaged using a 300 kV Titan Krios (Thermo Fisher Scientific) equipped with a GIF BioQuantum and K3 camera (Gatan). Movies were collected with Leginon^52^ with a pixel size of 1.0825 Å per px (micrograph dimensions of 5,760 × 4,092 px) over a defocus range of 0.8 to 1.8 μm with a 20 eV energy filter slit. Videos were recorded in counting mode (native K3 camera binning 2) with ∼25 e−/ Å^2^/s exposure rate in dose-fractionation mode with intermediate frames recorded every 40 ms over a 2 s exposure (50 frames per micrograph) to give an electron exposure of ∼51 e−/Å^2^. Dose-fractionated videos were gain-normalized, drift-corrected, summed and dose-weighted using MotionCor2^53^. The CTF was estimated for each summed image using the Patch CTF module in cryoSPARC v.3.3.2^54^. Particles were picked and extracted from the dose-weighted images with box size of 256 px using cryoSPARC blob picker and particle extraction. The entire dataset consisted of 18,565 motion-corrected images with 8,893,273 particles. Particles containing the RTC were sorted from junk particles using cryoSPARC heterogeneous refinement using a template of an RTC monomer^26^ and junk classes that pulled out particles containing the RTC as well as a volume for nsp9, not present in the template (Extended Data Fig. 4). This was followed by further particle curation using iterative heterogeneous refinement (n=5), resulting in 1,005,706 curated particles. These particles were refined using cryoSPARC local and global CTF refinements as well as non-uniform refinements (Extended Data Fig. 4). Particles were further processed through RELION v.3.1 Bayesian polishing^55^. Polished particles were then masked around the NiRAN domain and nsp9 then subtracted outside the mask (Extended Data Fig. 4). Subtracted particles were then classified using cryoSPARC v4.0.2 3D classification into 6 classes. The particles in these classes were reextracted to a box size of 256 px and processed through 2 rounds of RELION v.3.1 Bayesian polishing^55^. Polished particles were refined using cryoSPARC non-uniform refinements. The classes were then masked (Extended Data Fig. 4) around the NiRAN domain and nsp9 then subtracted outside the mask for cryoSPARC local refinements around the NiRAN domain active site. Locally refined maps were combined with the consensus maps into a composite map for each class using PHENIX ‘combine focused maps’ to aid model building^56^. Two classes were selected as the primary classes of interest for model building and analysis: 20mer-SL and GDP-Cap-i. These classes produced structures with following particle counts and nominal resolutions from cryoSPARC GSFSC: 20mer-SL (112,678 particles, 3.06) and GDP-Cap-i (113,000 particles, 3.01Å). Local resolution calculations were generated using blocres and blocfilt from the Bsoft package^57^ (Extended Data Figs. 5b, 5f). Angular distribution of particle orientations (Extended Data Figs. 5a, 5e) and directional resolution (Extended Data Figs. 5c, 5g), calculated through the 3DFSC package^58^ are shown for the final class. GSFSC (Extended Data Figs. 5b, 5f), calculated through cryoSPARC, and map-model FSC, calculated with Phenix mTriage, are shown for the final class (Extended Data Figs. 5d, 5h).

#### Model building and refinement

An initial model of the RTC+nsp9 was derived from Protein Data Bank (PDB) 7CYQ^25^. The models were manually fit into the cryo-EM density maps using Chimera^59^ and rigid-body and real-space refined using PHENIX real-space-refine^56^. For real-space refinement, rigid-body refinement was followed by all-atom and B factor refinement with Ramachandran and secondary structure restraints. Models were inspected and modified in Coot v.0.9.7^60^ and the refinement process was repeated iteratively.

## Data and code availability

All unique/stable reagents generated in this study are available without restriction from the lead contact, E.A.C. The cryo-EM density maps and atomic coordinates have been deposited in the EMDataBank and PDB as follows: UMP-i (EMD-40699, PDB 8SQ9), GDP-Cap-i (EMD-40708, PDB 8SQK), and 20mer-SL (EMD-40707, PDB 8SQJ). The model used for initial building (PDB 7CYQ) is available on the PDB.

## Acknowledgements

We thank D. Littler for plasmids and helpful discussions, and J. Perry for helpful discussions. Some of the work reported here was conducted at the Simons Electron Microscopy Center and the National Resource for Automated Molecular Microscopy and National Center for CryoEM Access and Training located at the NYSBC, supported by grants from the National Insitutes of Health (NIH) National Institute of General Medical Sciences (grant no. P41 GM103310), NYSTAR, the Simons Foundation (grant no. SF349247), the NIH Common Fund Transformative High Resolution Cryo-Electron Microscopy programme (grant no. U24 GM129539) and NY State Assembly Majority. Some of this work was conducted in the Rockefeller University Proteomics Resource Center and thus acknowledges funding from the Leona M. and Harry B. Helmsley Charitable Trust and Sohn Conferences Foundation for mass spectrometer instrumentation. This work was supported by NIH grant nos. P41 GM109824 and P41 GM103314 to B.T.C., and NIH grant no. R01 AI161278 (to E.A.C. and S.A.D.).

## Author contributions

G.S., O.F., J.C., A.B., Y.J.C., S.A.D., and E.A.C. conceived and designed this study. G.S., J.C., A.B., and Y.J.C. performed protein purification and biochemistry. O.F. provided critical reagents. P.D.B.O. and H.M. conducted mass spectrometry experiments. G.S. prepared cryo-EM specimens and processed all cryo-EM data. G.S., S.A.D., and E.A.C. built and analyzed atomic models. B.C., S.A.D., and E.A.C. supervised and acquired financial support. G.S. wrote the first draft of the manuscript; all authors contributed to the final version.

## Competing interests

All authors declare no competing interests.

## Notes

### Competing Interest Statement

The authors have declared no competing interest.

