## Supplementary Data for "Structural and functional insights into the enzymatic plasticity of the SARS-CoV-2 NiRAN Domain"

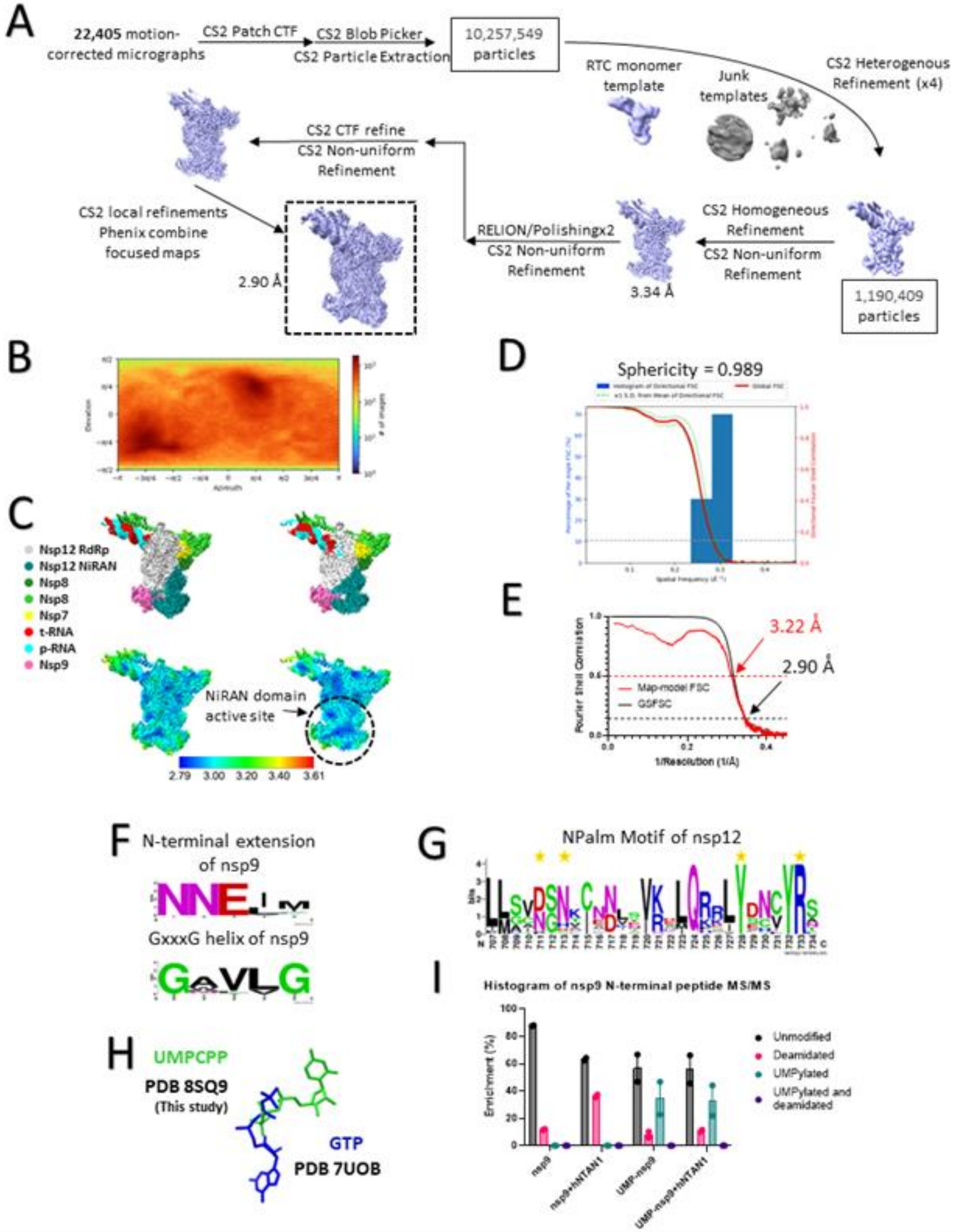
A
22,405 motion-corrected micrographs
CS2 Patch CTF
CS2 Blob Picker
CS2 Particle Extraction
10,257,549 particles
RTC monomer template
Junk templates
CS2 Heterogenous Refinement (x4)
CS2 CTF refine
CS2 Non-uniform Refinement
CS2 local refinements
Phenix combine focused maps
2.90 Å
RELION/Polishingx2
CS2 Non-uniform Refinement
3.34 Å
CS2 Homogeneous Refinement
CS2 Non-uniform Refinement
1,190,409 particles
B
D
Sphericity = 0.989
C
Nsp12 RdRp
Nsp12 NiRAN
Nsp8
Nsp8
Nsp7
t-RNA
p-RNA
Nsp9
NiRAN domain active site
2.79 3.00 3.20 3.40 3.61
E
3.22 Å
2.90 Å
Map-model FSC
GSFSC
Fourier Shell Correlation
1/Resolution (1/Å)
F
N-terminal extension of nsp9
GxxxG helix of nsp9
G
NPalm Motif of nsp12
H
UMPCPP
PDB 8SQ9
(This study)
GTP
PDB 7UOB
I
Histogram of nsp9 N-terminal peptide MS/MS
Enrichment (%)
Unmodified
Deamidated
UMPylated
UMPylated and deamidated
nsp9
nsp9+hNTAN1
UMP-nsp9
UMP-nsp9+hNTAN1

**Figure S1. Cryo-EM processing pipeline and analysis for UMP-I and functional analysis of nsp9 NMPylation**

(A) Cryo-EM processing pipeline for the dataset producing the UMP-i map.

(B) Angular distribution plot for the 20mer-SL dataset, calculated in cryoSPARC. Scale depicts number of particles assigned to a specific angular bin.

(C) Nominal 3.06 Å resolution cryo-EM reconstruction filtered by local resolution and colored according to fitted model chain above the same cryo-EM reconstruction colored by local resolution. Right panels are clipped to reveal NiRAN active site.

(D) Directional 3D FSC for 20mer-SL, determined with 3DFSC.

(E) Plot containing the gold-standard FSC (GSFSC) and the model-map FSC for the 20mer-SL dataset. GSFSC calculated by comparing two half maps from cryoSPARC and model-map FSC calculated with Phenix Mtriage. The dotted lines represent the 0.5 and 0.143 FSC cutoffs.

(F) UMPCPP from this study and GTP (PDB = 7UOB) superposed to show clashing of phosphates and mutually exclusive positioning of GTP binding in a 'base in' or 'base up' pose.

(G) Sequence conservation logos of the N terminal extension and the GxxxG helix in α and β coronavirus nsp9s.

(H) Sequence conservation logos of the NPalm motif in α and β coronavirus nsp12s. Residues that make polar contacts with nsp9, UMPCPP, or RNA-nsp9 in UMP-I and GDP-Cap-i are starred.

(I) Histogram showing the N-terminal peptide enrichment from MS/MS of hNTAN1 deamidation reactions based on the N-terminal peptide modification status. Results are the average of two independent experiments and errors represent the SEM.

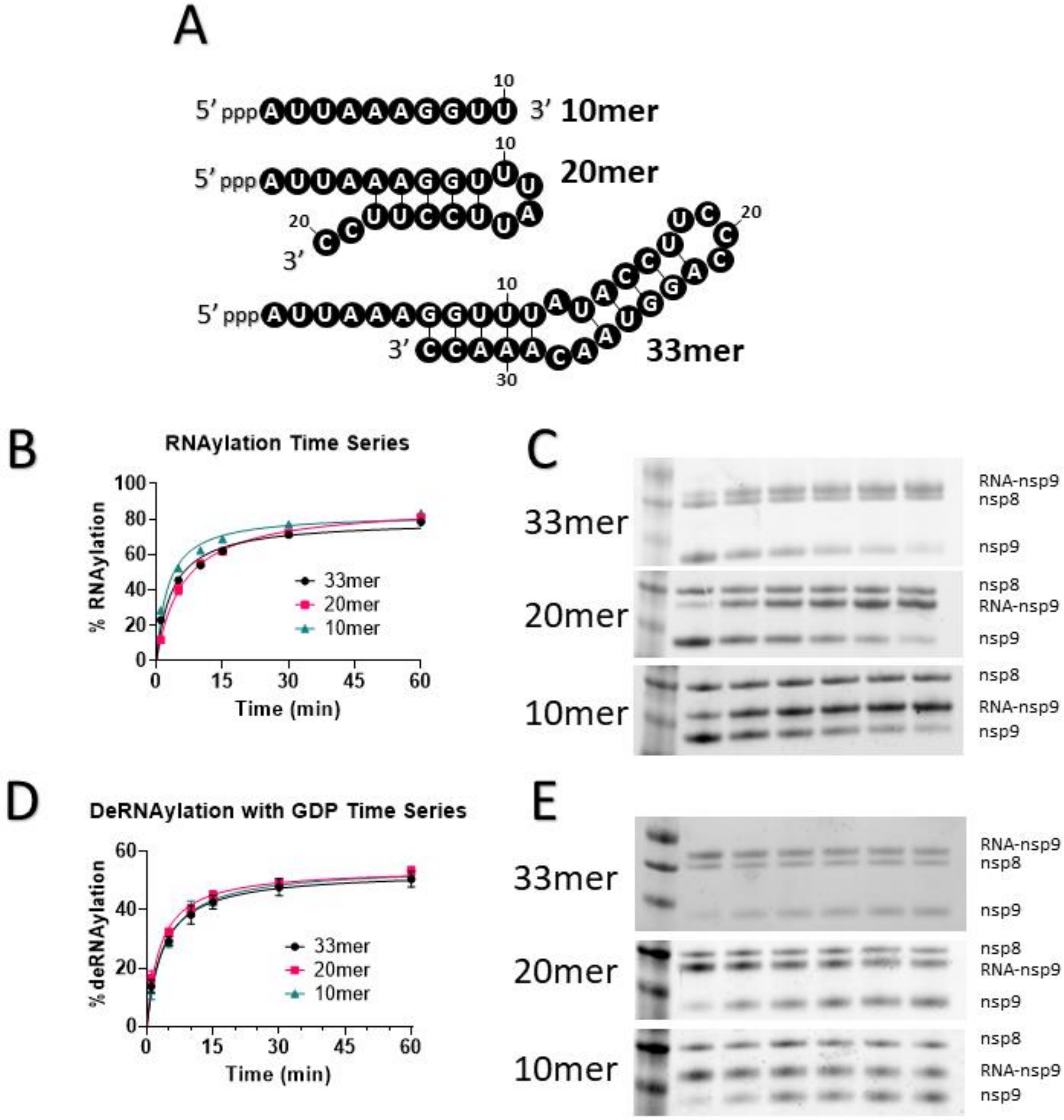


**Figure S2. Biochemical analysis of NiRAN domain oligo substrate preferences**

(A) The three different pppRNA constructs used in this study are shown with their predicted secondary structures[61].

(B) NiRAN domain mediated RNAylation of nsp9 time series with three different pppRNAs. Reactions were quenched in 4X LDS + 50 mM DTT, run on SDS-PAGE, and then bands were quantified using gel densitometry. Numbers used for analysis are the intensity of the RNA-nsp9 band divided by the sum of the RNA-nsp9 and nsp9 band intensities. These are the results of 3 independent experiments.

(C) Representative gels for the RNAylation experiments shown in (B).

(D) NiRAN domain mediated deRNAylation with GDP of RNA-nsp9 time series with three different RNA-nsp9s. Samples were analyzed as in (B). These are the results of 3 independent experiments.

(E) Representative gels for the deRNAylation experiments shown in (D).

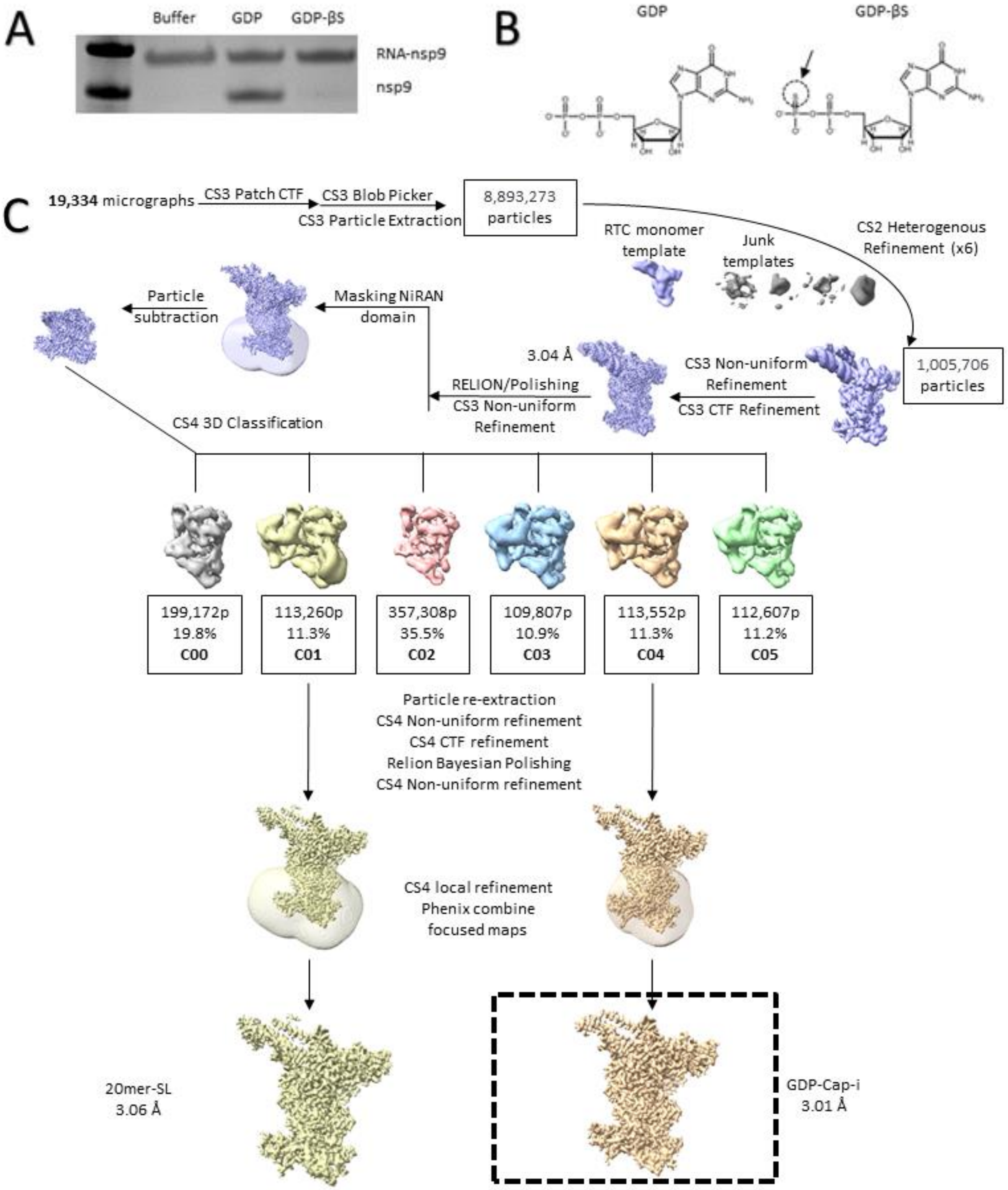


**Figure S3. Cryo-EM processing pipeline for dataset producing GDP-Cap-i and 20mer SL**
(A) Gel demonstrating that GDP-βS is unreactive for deRNAylation.
(B) Chemical structure comparison between GDP and GDP- βS.
(C) Cryo-EM processing pipeline for the dataset producing the GDP-Cap-i and 20mer-SL maps.

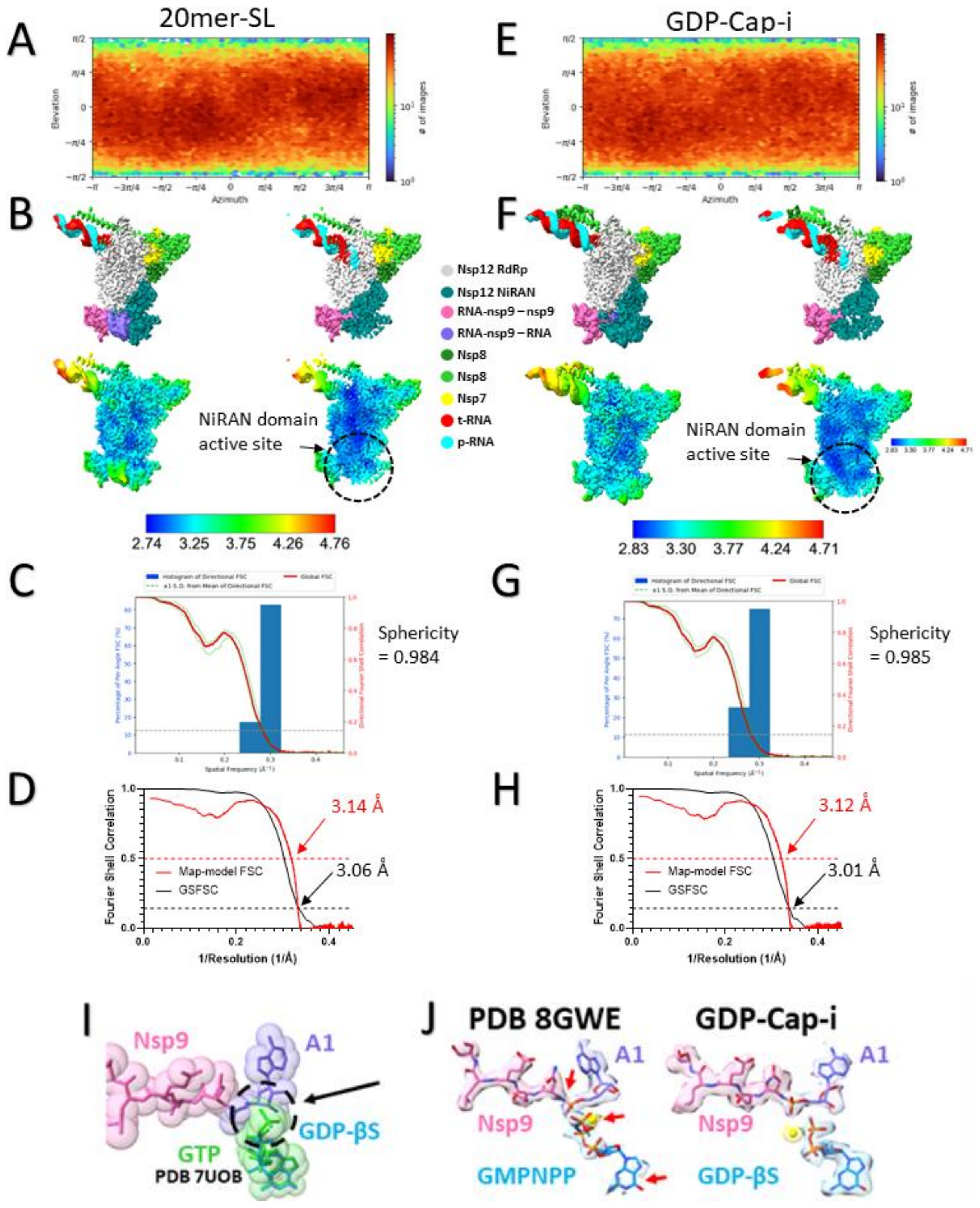


**Figure S4. Cryo-EM analysis for GDP-Cap-i and 20mer-SL maps and comparison to other published models**

(A) Angular distribution plot for the 20mer-SL dataset, calculated in cryoSPARC. Scale depicts number of particles assigned to a specific angular bin.
(B) Nominal 3.06 Å resolution cryo-EM reconstruction filtered by local resolution and colored according to fitted model chain above the same cryo-EM reconstruction colored by local resolution. Right panels are clipped to reveal NiRAN active site.
(C) Directional 3D FSC for 20mer-SL, determined with 3DFSC.
(D) Plot containing the gold-standard FSC (GSFSC) and the model-map FSC for the 20mer-SL dataset. GSFSC calculated by comparing two half maps from cryoSPARC and model-map FSC calculated with Phenix Mtriage. The dotted lines represent the 0.5 and 0.143 FSC cutoffs.
(E) Angular distribution plot for the GDP-Cap-i dataset, calculated in cryoSPARC. Scale depicts number of particles assigned to a specific angular bin.
(F) Nominal 3.01 Å resolution cryo-EM reconstruction filtered by local resolution and colored according to fitted model chain above the same cryo-EM reconstruction colored by local resolution. Right panels are clipped to reveal NiRAN active site.
(G) Directional 3D FSC for GDP-Cap-i, determined with 3DFSC.
(H) Plot containing the gold-standard FSC (GSFSC) and the model-map FSC for the GDP-Cap-i dataset. GSFSC calculated by comparing two half maps from cryoSPARC and model-map FSC calculated with Phenix Mtriage. The dotted lines represent the 0.5 and 0.143 FSC cutoffs.
(I) GDP-Cap-i aligned with GTP from 7UOB with an arrow indicating that GTP would clash with the RNA-nsp9 in GDP-Cap-i.
(J) Comparison of RNA-nsp9 and GDP-βS/GMPNPP from GDP-Cap-i and PDB 8GWE with the red arrows pointing out the following issues with the modeling in PDB 8GWE. A lack of map density to support the modeling of a GMPNPP in the NiRAN active site. The Mg2+ is modeled without map density and clashes with the GMPNPP. Negatively charged GMPNPP phosphate groups are modeled closely to the negatively charged phosphoramidate. These issues with modeling and with the absence of data make the reported structure invalid for analysis of CoV mRNA capping.

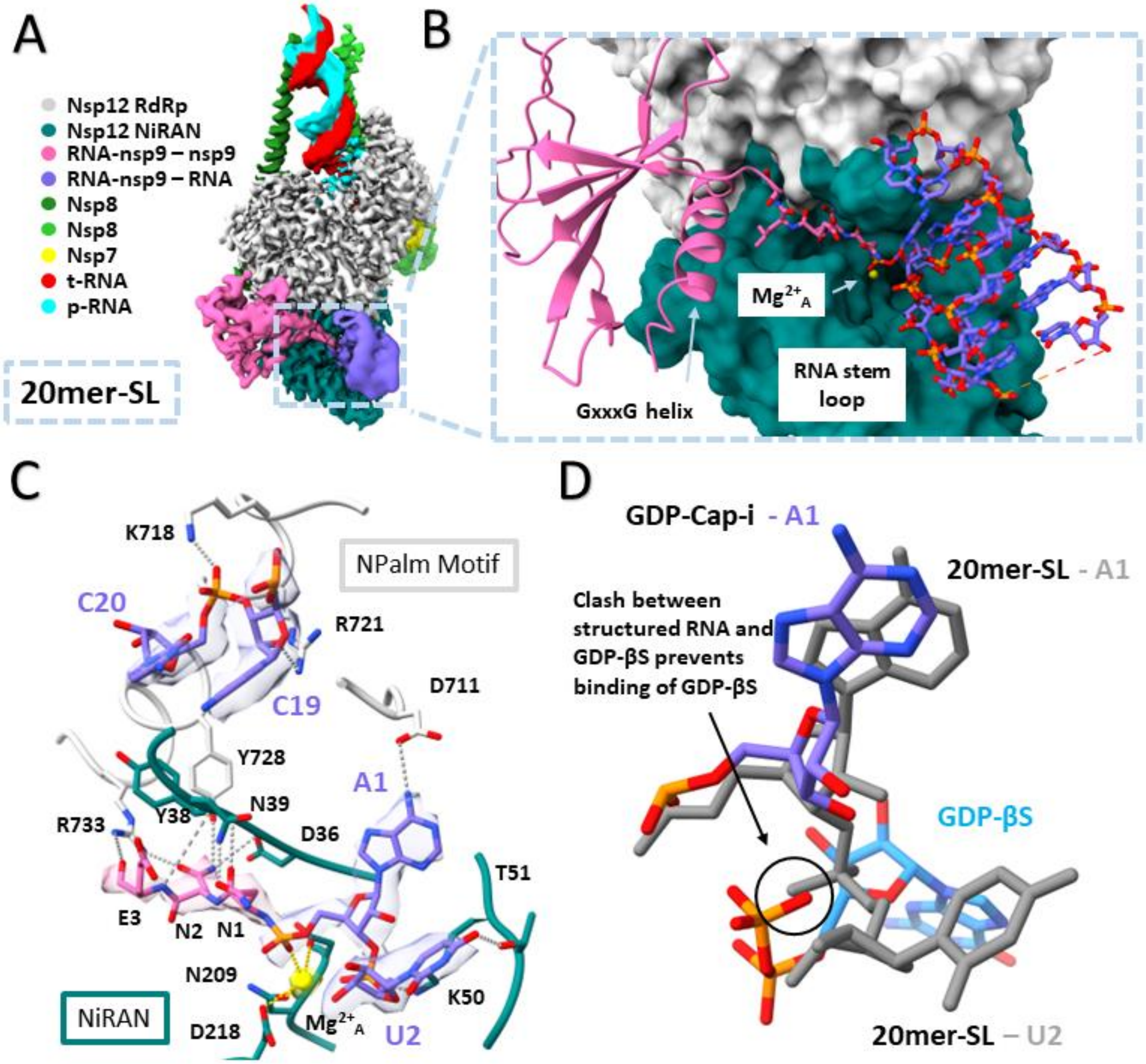


**Figure S5. Structural analysis of a catalytically incompetent RNA-nsp9 binding mode**

(A) Locally filtered cryo-EM map of the 3.0 Å resolution structure reveals the general architecture of a trapped catalytically incompetent binding mode of 20mer RNA-nsp9 to the RTC. Map is colored according to fitted model chains.

(B)Close up of the NiRAN domain active site. Nsp12 is portrayed as a surface and is colored according to domain.

(C) Close up of the interface between RNA-nsp9's N-terminal extension and the RNA stem loop with nsp12. Interacting residues are shown as sticks. RNA-nsp9 is shown within locally filtered cryo-EM map and polar interactions are shown with dashed gray lines.
(D) A1 and GDP- βS from C04 is superposed with A1 and U2 from C01 demonstrating a clash between GDP- βS and the phosphodiester bond between A1 and U2 that prevents binding of GDP- βS in C01.

Table S1. Cryo-EM data collection, refinement, and validation statistics.

| | UMP-i (EMDB-40699) (PDB 8SQ9) | GDP-Cap-i (EMDB- 40708) (PDB 8SQK) | 20mer-SL (EMDB- 40707) (PDB 8SQJ) |
|---|---|---|---|
| ***Data collection and processing*** | | | |
| Magnification | 81,000 | 81,000 | 81,000 |
| Voltage (kV) | 300 | 300 | 300 |
| Electron exposure (e–/Å$^2$) | 51.18 | 51.51 | 51.51 |
| Defocus range (μm) | 0.7 μm to 2.5 μm | 0.8 μm to 1.8 μm | 0.8 μm to 1.8 μm |
| Pixel size (Å) | 1.069 | 1.085 | 1.085 |
| Symmetry imposed | C1 | C1 | C1 |
| Initial particle images (no.) | 10,257,549 | 8,893,273 | 8,893,273 |
| Final particle images (no.) | 1,190,409 | 113,552 | 113,260 |
| Map resolution (Å) FSC threshold = 0.143 | 2.90 | 3.06 | 3.01 |
| Map resolution range (Å) | 2.79-3.61 | 2.83-4.71 | 2.74-4.76 |
| ***Refinement*** | | | |
| *Initial model used (PDB code)* | 7CYQ | 8SQ9 | 8SQ9 |
| Model resolution (Å) FSC threshold = 0.5 | 3.22 | 3.12 | 3.14 |
| Model composition | | | |
| Non-hydrogen atoms | 13,325 | 13,354 | 13,668 |
| Protein residues | 1,469 | 1,487 | 1486 |
| Nucleic acid residues | 67 | 70 | 85 |
| Ligands | 6 | 3 | 3 |
| B factors (Å$^2$) | | | |
| Protein | 90.55 | 55.17 | 59.19 |
| Nucleic acids | 189.51 | 150.68 | 150.75 |
| Ligand | 93.79 | 80.24 | 68.51 |
| R.m.s. deviations | | | |
| Bond lengths (Å) | 0.004 | 0.003 | 0.004 |
| Bond angles (°) | 0.482 | 0.543 | 0.500 |
| Validation | | | |
| MolProbity score | 1.83 | 1.71 | 1.84 |
| Clashscore | 4.41 | 4.20 | 3.47 |
| Poor rotamers (%) | 2.38 | 1.97 | 3.69 |
| Ramachandran plot | | | |
| Favored (%) | 95.26 | 95.87 | 96.07 |
| Allowed (%) | 4.67 | 4.13 | 3.93 |
| Disallowed (%) | 0.07 | 0 | 0 |

**Table S2. RNA constructs used for cryo-EM analyses.**

| Structure | RNA Scaffolds |
|---|---|
| UMP-i | $^{5'}$cgcguagcaugcuacGUCAUUCUCCacgcgaagcA$^{3'}$<br>$^{3'}$gcgcaucguacgaugCAGUAAGAGGugcgcuucguAcuguuguuuUACCCCUAUC$^{5'}$ |
| GDP-Cap-I | $^{5'}$cgcguagcaugcuacGUCAUUCUCCacgcgaagcaU$^{3'}$<br>$^{3'}$gcgcaucguacgaugCAGUAAGAGGugcgcuucguAcuguuguuuUACCCCUAUC$^{5'}$ |
| 20mer-SL | $^{5'}$cgcguagcaugcuacGUCAUUCUCCacgcgaagcaU$^{3'}$<br>$^{3'}$gcgcaucguacgaugCAGUAAGAGGugcgcuucguAcuguuguuuUACCCCUAUC$^{5'}$ |

**Table S3. Coronavirus species and references used for nsp9 amino acid sequence alignment.**

| Virus | Reference |
|---|---|
| Betacoronavirus HKU24 | >YP_009113022.1 |
| Human coronavirus OC43 | >YP_009555252.1 |
| Bovine coronavirus | >NP_150073.3 |
| Human coronavirus HKU1 | >YP_173236.1 |
| Murine hepatitis virus strain JHM | >YP_209229.2 |
| Rat coronavirus Parker | >YP_003029844.1 |
| Murine hepatitis virus | >YP_009915696.1 |
| Betacoronavirus Erinaceus/VMC/DEU/2012 | >YP_009513008.1 |
| Middle East respiratory syndrome-related coronavirus | >YP_009047202.1 |
| Tylonycteris bat coronavirus HKU4 | >YP_001039952.1 |
| Pipistrellus bat coronavirus HKU5 | >YP_001039961.1 |
| Rousettus bat coronavirus HKU9 | >YP_001039970.1 |
| Rousettus bat coronavirus | >YP_009273004.1 |
| Bat Hp-betacoronavirus/Zhejiang2013 | >YP_009072438.1 |
| Severe acute respiratory syndrome coronavirus 2 | >YP_009724389.1 |
| SARS coronavirus Tor2 | >NP_828849.7 |
| Bat coronavirus BM48-31/BGR/2008 | >YP_003858583.1 |
| Wencheng Sm shrew coronavirus | >YP_009824973.1 |
| Bat coronavirus CDPHE15/USA/2006 | >YP_008439200.1 |
| Scotophilus bat coronavirus 512 | >YP_001351683.1 |
| Porcine epidemic diarrhea virus | >NP_598309.2 |
| Bat coronavirus 1A | >YP_001718603.1 |
| Miniopterus bat coronavirus HKU8 | >YP_001718610.1 |
| BtMr-AlphaCoV/SAX2011 | >YP_009199608.1 |
| Alphacoronavirus Bat-CoV/P.kuhlii/Italy/3398-19/2015 | >YP_009755889.1 |
| BtNv-AlphaCoV/SC2013 | >YP_009201729.1 |
| BtRf-AlphaCoV/HuB2013 | >YP_009199789.1 |
| Rousettus bat coronavirus HKU10 | >YP_006908641.2 |
| BtRf-AlphaCoV/YN2012 | >YP_009200734.1 |
| Rhinolophus bat coronavirus HKU2 | >YP_001552234.1 |
| Camel alphacoronavirus | >YP_009194637.1 |
| Human coronavirus 229E | >NP_073549.1 |
| NL63-related bat coronavirus | >YP_009824965.1 |
| Human coronavirus NL63 | >YP_003766.2 |
| NL63-related bat coronavirus | >YP_009328933.1 |
| Lucheng Rn rat coronavirus | >YP_009336483.1 |
| Coronavirus AcCoV-JC34 | >YP_009380519.1 |
| Feline infectious peritonitis virus | >YP_004070193.2 |
| Swine enteric coronavirus | >YP_009199240.1 |
| Transmissible gastroenteritis virus | >NP_058422.1 |
| Ferret coronavirus | >YP_009256195.1 |
